# Perturb-tracing enables high-content screening of multiscale 3D genome regulators

**DOI:** 10.1101/2023.01.31.525983

**Authors:** Yubao Cheng, Mengwei Hu, Bing Yang, Tyler B Jensen, Tianqi Yang, Ruihuan Yu, Zhaoxia Ma, Jonathan S D Radda, Shengyan Jin, Chongzhi Zang, Siyuan Wang

## Abstract

Three-dimensional (3D) genome organization becomes altered during development, aging, and disease^1–23^, but the factors regulating chromatin topology are incompletely understood and currently no technology can efficiently screen for new regulators of multiscale chromatin organization. Here, we developed an image-based high-content screening platform (Perturb-tracing) that combines pooled CRISPR screen, a new cellular barcode readout method (BARC-FISH), and chromatin tracing. We performed a loss-of-function screen in human cells, and visualized alterations to their genome organization from 13,000 imaging target-perturbation combinations, alongside perturbation-paired barcode readout in the same single cells. Using 1.4 million 3D positions along chromosome traces, we discovered tens of new regulators of chromatin folding at different length scales, ranging from chromatin domains and compartments to chromosome territory. A subset of the regulators exhibited 3D genome effects associated with loop-extrusion and A-B compartmentalization mechanisms, while others were largely unrelated to these known 3D genome mechanisms. We found that the ATP-dependent helicase CHD7, the loss of which causes the congenital neural crest syndrome CHARGE^24^ and a chromatin remodeler previously shown to promote local chromatin openness^25–27^, counter-intuitively compacts chromatin over long range in different genomic contexts and cell backgrounds including neural crest cells, and globally represses gene expression. The DNA compaction effect of CHD7 is independent of its chromatin remodeling activity and does not require other protein partners. Finally, we identified new regulators of nuclear architectures and found a functional link between chromatin compaction and nuclear shape. Altogether, our method enables scalable, high-content identification of chromatin and nuclear topology regulators that will stimulate new insights into the 3D genome functions, such as global gene and nuclear regulation, in health and disease.

## Main

The spatial organization of chromatin is linked to many genomic functions and shows intriguing dynamics in a variety of biological processes and diseases^1–23^. Chromatin organization occurs at many levels and length scales, from local accessibility to inter-domain contacts to global nuclear architecture. Studies on effectors such as histone modification enzymes or chromatin loop organizers *e.g.* CTCF have bolstered our understanding of chromatin architecture and provided new drug targets for a wide range of diseases, but our overarching understanding of how chromatin is regulated across length scales and in different cell types and conditions remains limited. It is a significant challenge to systematically identify new molecular regulators of complex three-dimensional (3D) genome architecture and form testable hypotheses about their mechanisms of action. Important 3D genome regulators have been primarily discovered by perturbing one candidate gene at a time^28–30^ or by plate-based screening focusing on relatively low-content phenotypes such as the spatial distance between one pair of genomic loci^31, 32^. Recently, high-content CRISPR screens combining Perturb-seq and single-cell ATAC-seq technologies have enabled the high-throughput discovery of candidate regulators of local chromatin accessibility^33, 34^. However, we still lack scalable, broadly applicable methods to efficiently screen for regulators of higher-order 3D chromatin folding architectures, especially at the length scales of topologically-associating domains (TADs, also known as contact domains), chromatin compartments, and chromosome territories^35–41^.

TADs largely confine the scope of promoter-enhancer interactions^42, 43^ and are structural units of chromatin with different DNA replication timing^44^ and mutation susceptibility^45^. TADs are further sorted into segregated A (active) and B (inactive) compartments in each chromosome territory^36^. Whole chromosome territory compaction has been observed in X chromosome inactivation^46^ and cellular senescence^47^. Defining the regulatory landscape and architectural basis of chromatin folding at the length scales that are relevant to each genomic feature is critical to understanding their functions and dynamics in development, aging, and disease^3, 9, 48^. Assessing multiple length scales in parallel is technically challenging and currently only possible with single-gene perturbations.

We endeavored to develop a technique that would allow for discovery of new factors influencing chromatin architecture as well as providing clues into which length scales and chromatin features they act on. To this end, here we present a new method termed Perturb-tracing that combines the power of pooled CRISPR screening of candidate regulators with high-content readout of chromatin organization over multiple length scales. Our high-throughput, high-content genetic perturbation screen is combined with super-resolved *in situ* tracing of complex chromatin folding conformations and imaging of nuclear architectures across multiple length scales in human cells. A key innovation of our technology is the decoding technique we devised, termed BARC-FISH, which enzymatically amplifies the barcode of each sgRNA *in situ* for robust decoding using FISH, which is compatible with the multi-scale 3D chromatin mapping required to assess high-content phenotypic readouts for each barcode. After validating our new method, we applied it to screen 137 candidate genes with 420 single guide RNAs (sgRNAs) and imaged 30 chromatin or cellular targets per cell, generating 12,600 imaging target-perturbation combinations. The screen identified 26 top hit novel regulators of 3D genome organization at different length scales. Correlation analyses revealed regulators working in conjunction with known 3D genome regulatory mechanisms. In particular, we identified CHD7, a chromatin remodeler critical for normal neural crest development, as a long-range chromatin compactor, which is unexpected from its known activity in promoting local chromatin accessibility. We found that CHD7 compacts chromatin over long range independently of its chromatin remodeling function and known interaction partners, and globally suppresses gene expression. *In vitro* reconstitution further proved that CHD7 directly compacts and condenses DNA in an ATP-independent manner. In addition, we discovered a general link between chromatin compaction regulation and the maintenance of nuclear sphericity, and showed that CHD7 depletion leads to a multi-lobed nuclear shape. We interpreted this linkage with polymer simulation proposing that chromatin folding regulates nuclear morphology. These results offer new hypotheses on the molecular and cellular mechanisms of complex diseases associated with CHD family proteins and other regulators. Altogether, our work presented here provides a new platform for the discovery and characterization of new chromatin regulators across multiple length scales and paves the way to build a global map of how 3D genomic architecture is regulated in diverse contexts.

### Perturb-tracing enables image-based pooled CRISPR screen of chromatin and nuclear organization regulators

To systematically discover novel regulators of chromatin conformation, we developed an image-based method termed Perturb-tracing to screen chromatin structures in individual knockout cells. Briefly, we used CRISPR-Cas9 technology to generate a pooled library of knockout A549 human lung cancer cells that co-express Cas9 with a single guide RNA (sgRNA) and a unique RNA barcode (Fig. 1a and Extended Data Fig. 1a,b). The barcode RNA comprised ten regions, each encoding one of three sequences (Fig. 1b), a design inspired by a recent report^49^. Each region is analogous to a ternary digit (with values “0”, “1”, or “2”) in computation, and each ten-digit barcode was uniquely paired with an sgRNA in the same cell (Fig. 1a,b). We developed “Barcode Amplification by Rolling Circle and Fluorescence *In Situ* Hybridization” (BARC-FISH) to visually read out the barcode RNAs in single cells, and thereby identify the genetic perturbation in each cell (see below). To map the phenotypic effects of each knockout on 3D spatial organization of numerous genomic loci in single cells, we performed the highly multiplexed DNA FISH method known as chromatin tracing^50^ on chromosome 22 (chr22) (Fig. 1a). Finally, we employed a high-throughput computational approach to identify conserved features and systematic changes of chromatin organization caused by the same genetic perturbation in multiple cells, for every gene knockout in the pooled screen.

**Fig. 1.**
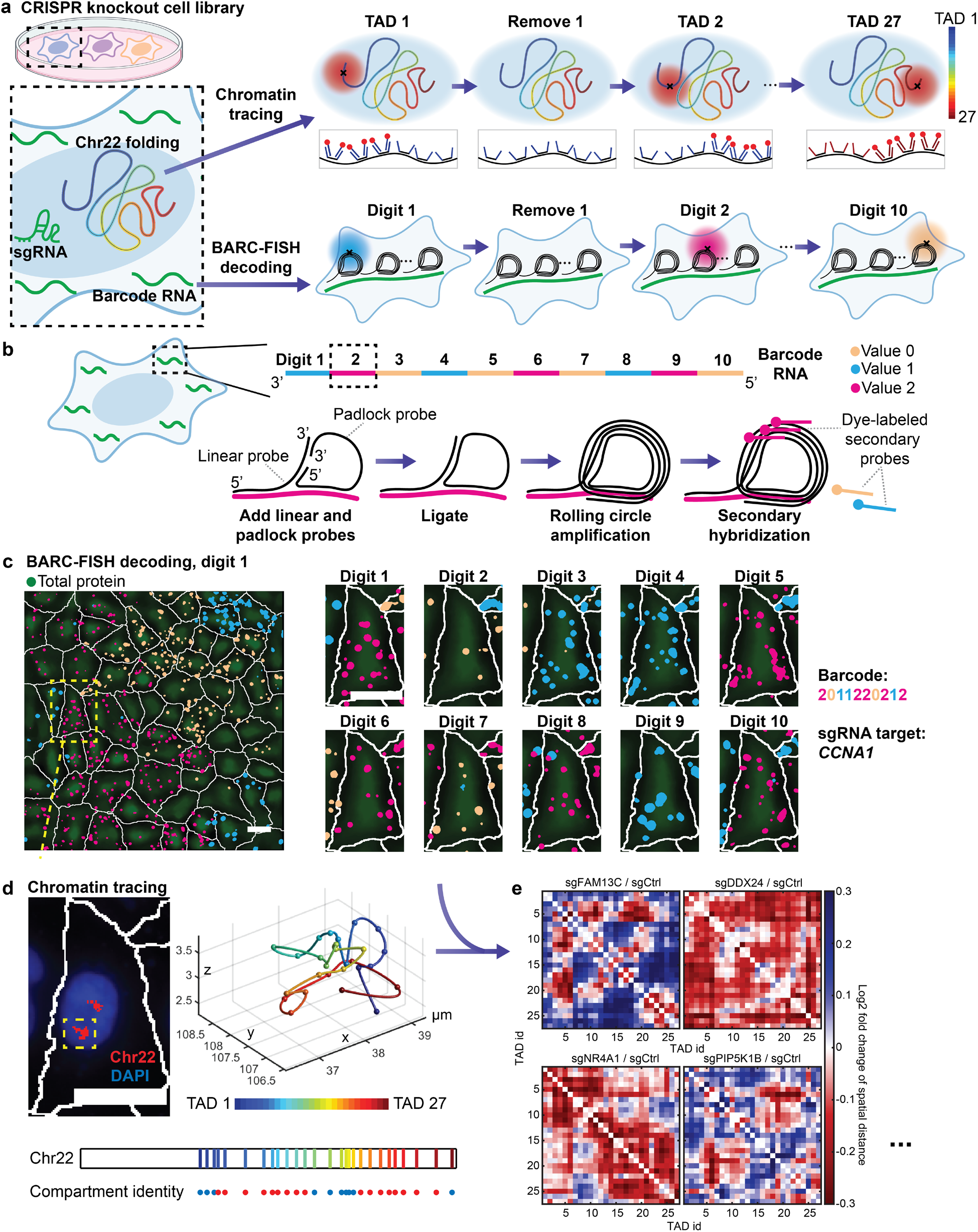
Perturb-tracing enables image-based pooled CRISPR screen of chromatin and nuclear organization regulators. **a**, Schematic of the screening approach. A lentivirus library encoding paired sgRNAs and barcode RNAs was transduced into human A549 cells expressing Cas9 protein. The organization of chr22 was determined by chromatin tracing and the identity of the knockout gene was determined by BARC-FISH decoding of the barcode RNAs. For chromatin tracing, all 27 TADs spanning chr22 were sequentially visualized in a multiplexed DNA FISH procedure. For BARC-FISH decoding, 10 digits of the barcode were amplified and sequentially imaged. **b**, A scheme of the BARC-FISH method. In each cell, the expressed barcode RNA was composed of 10 “digits”, and each digit had one of three different values (values 0, 1 and 2, represented by orange, cyan, and magenta respectively). Each digit was hybridized with a linear probe and padlock probe, enabling ligation of the padlock probe, which was then subjected to rolling circle amplification, generating an amplicon containing multiple copies of the digit sequences. Dye-labelled secondary probes were then introduced for imaging, reading out the value of the digit. **c**, An example of BARC-FISH decoding. Left: A representative field of view from the screen with BARC-FISH signals shown in orange, cyan, and magenta, cell segmentation shown as white lines, and total protein stain in green. Right: The yellow-boxed cell in the left panel in 10 rounds of decoding. Scale bars: 20 µm. **d**, Chromatin tracing of the yellow-boxed cell in **c**. Left: An image of the cell, with the traces of the two copies of chr22 shown in red and DAPI stain shown in blue. Right: 3D chromatin trace of the yellow-boxed chromosome in the left panel. The 3D positions of each TAD were shown as pseudo-colored spots, connected with a smooth curve. Below: The genomic positions of TADs 1-27 on chr22, and their corresponding compartment identity (red: compartment A; blue: compartment B). Scale bar: 20 µm. **e**, Example matrices of log2 fold changes of inter-TAD spatial distances for selected hits from the screen.

For BARC-FISH, we adopted a high-efficiency rolling circle amplification (RCA) strategy from an *in situ* sequencing method^51^. Specifically, a linear probe and a “padlock” probe are hybridized to each “digit” in the RNA barcode. Circularization and ligation of the padlock probe generate a template for RCA, primed by the linear probe, which in turn locally generates many copies of part of the digit sequence (Fig. 1b). For each of the ten digits, we hybridized dye-labeled secondary probes to the RCA product and observed a strong signal over background (Fig. 1b, c). To detect the value of each digit in individual cells, we sequentially applied three-color secondary probes for each digit, and imaged the fluorescence signals from the labeled RCA products over ten rounds of three-color sequential FISH imaging (Fig. 1c). After each round of imaging, the fluorescence signals were removed from the cells before the next round of sequential FISH. Finally, the fluorescence signals from all rounds of imaging were computationally converted to barcode values, and subsequently to sgRNA identities for individual cells by referring to the barcode-sgRNA associations mapped by high-throughput sequencing (Methods). With high signal-to-background ratio and an error-correcting decoding algorithm (Methods), BARC-FISH achieved robust decoding while being compatible with chromatin tracing (Fig. 1c,d and Extended Data Fig. 1c,d), allowing us to match sgRNA identities underlying genetic perturbations to 3D genome phenotypes (Fig. 1e).

For our screen, we generated a plasmid library of 420 sgRNAs composed of 10 non-targeting control sgRNAs and 410 sgRNAs targeting 137 selected genes (a coverage of 2-3 sgRNAs per target gene) (Supplementary Table 1). The plasmid library was cloned using a high-throughput pooled cloning strategy (Extended Data Fig. 2a, Methods). This strategy randomly paired the 10-digit barcodes with the sgRNAs, and used a bottlenecking method to ensure unique mapping of each barcode to a single sgRNA. With this strategy, up to ∼5,000 sgRNAs can be distinguished with the current barcoding scheme (Methods). It is possible to extend the barcode sequence to allow for more perturbations in a single screen. The 137 selected genes included known chromatin conformation regulators such as NIPBL and CTCF as positive controls^10^, and primarily genes encoding nuclear proteins that are differentially expressed by more than five-fold upon oncogene-induced senescence^52^, given the extensive 3D genome reorganization during this process^47, 48^. We generated the cell knockout library by lentiviral transduction of the plasmid library and puromycin selection of the transduced cells, and detected 8 non-targeting sgRNA controls and 404 sgRNAs corresponding to all 137 targeted genes, suggesting an sgRNA dropout rate of 1.9% with our library construction procedure. To estimate the knockout efficiency in our cell background, we transduced A549-Cas9 cells with selected individual sgRNA constructs and performed next-generation sequencing on the target genomic DNA of the polyclonal cells after transduction and selection. The results showed knockout efficiencies of 43-70% for the selected sgRNAs (Extended Data Fig. 2b), close to the predicted knockout efficiencies in previously reported CRISPR editing prediction model^53, 54^. This less-than-100% knockout efficiency is intrinsic to CRISPR and is expected to potentially lead to a mixture of wildtype and knockout phenotypes in subpopulations of cells carrying the same sgRNA, but we expect to still capture knockout phenotypes that are sufficiently strong in population analyses.

### Identification of regulators of multi-scale chromatin folding

For chromatin tracing, we mapped the conformation of chr22 as a model (due to its short genomic length and the lack of known structural variations on chr22 in A549 cells) at the TAD-to-chromosome length scales by pinpointing the central 100-kb regions of all 27 TADs spanning chr22 (Fig. 1d). To exclude the potential influence of cell cycle on chromatin conformation, we incorporated Geminin staining and only included G1 phase cells in our analyses (Extended Data Fig. 2c)^55^. Individual G1 phase A549-Cas9 cells contained 2-4 chr22 traces each. We analyzed 57,286 traces containing 1,407,797 3D positions from 17,304 cells.

To identify regulators of 3D genome organization, first we investigated the spatial distances between adjacent TADs on chr22. Consistent with recent imaging studies focusing on single pairs of adjacent TADs^56, 57^, we found that loss of the known cohesin loader NIPBL led to a significant increase in adjacent TAD distance, whereas knocking out CTCF significantly decreased adjacent TAD distance as expected^56, 57^ (Fig. 2a,b). These positive control results are consistent with the opposing roles of CTCF and NIPBL in loop extrusion^28–30, 56–59^, and cross-validated our screening method. Importantly, other knockout hits also altered the adjacent TAD distances, revealing new candidate regulators of chromatin organization and providing hints as to the length scale at which they operate. We observed that knocking out the tumor suppressor RB1, MRVI1 and PIP5K1B increased the adjacent TAD distance, while knocking out GLDC, the nuclear receptor NR4A1 and ZNF114 caused the opposite phenotype (Fig. 2a-c).

**Fig. 2.**
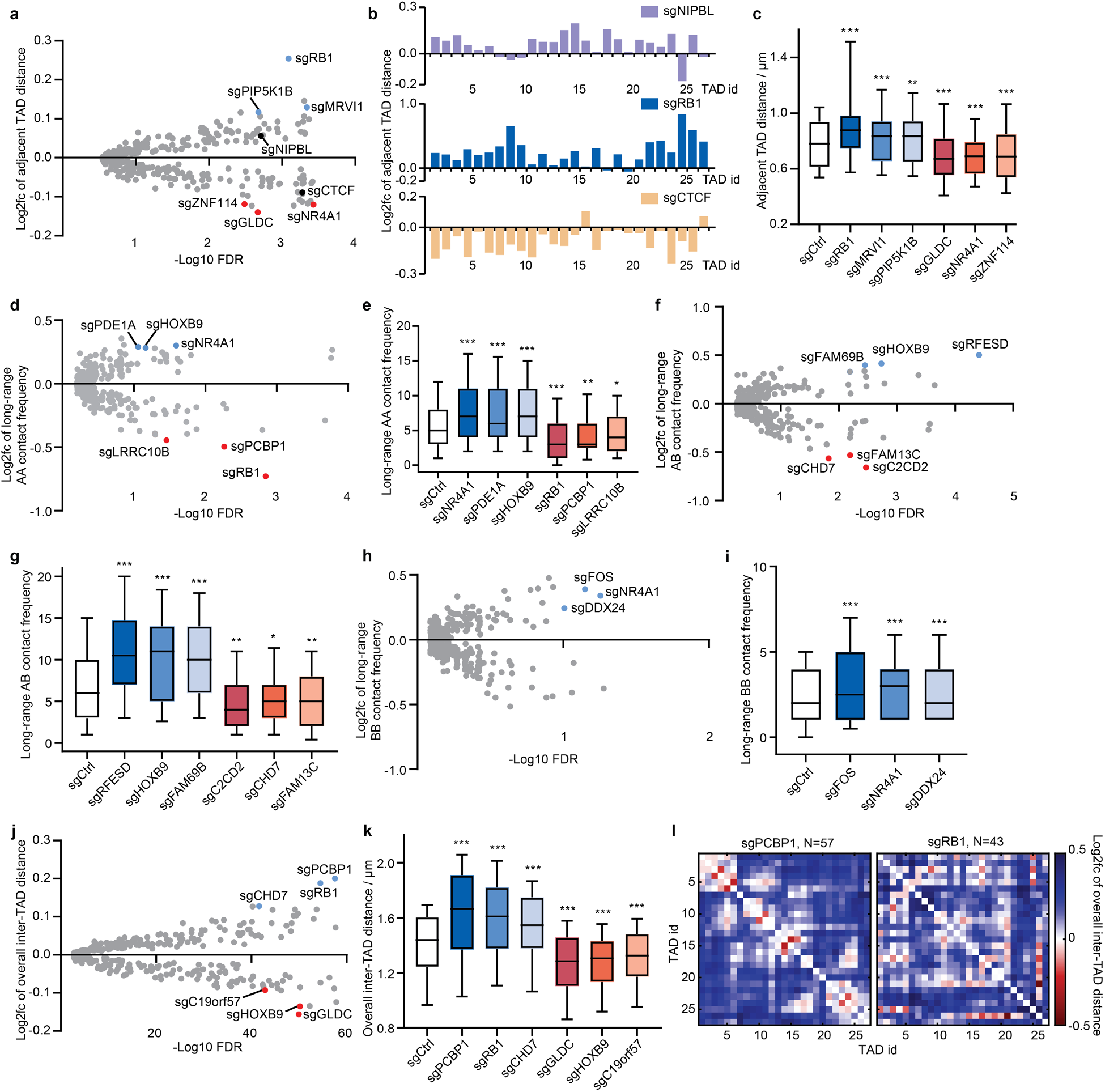
Perturb-tracing screen identified regulators of multi-scale chromatin folding. **a**, Log2 fold change (log2fc) of spatial distance between adjacent TADs versus -log10 false discovery rate (FDR) for each perturbation. Each dot represents a perturbation in the screen library. In all volcano plots, the top hits (nuclear proteins with the largest log2fc and FDRs<0.1) in both directions are indicated with blue (knockout leads to upregulation) and red dots (knockout leads to downregulation), respectively. The top candidate genes which when knocked out led to increased adjacent TAD distances are: RB1, MRVI1 and PIP5K1B; the top candidate genes which when knocked out caused decreased adjacent TAD distances are: GLDC, NR4A1 and ZNF114. Positive controls (NIPBL and CTCF) are marked in black. **b**, Log2 fold change of adjacent TAD distance across chr22 for selected hits. **c**, Spatial distances between adjacent TADs for non-targeting control and selected hits. **d**, Log2 fold change of long-range A-A contact frequency versus -log10 FDR for each perturbation. Top three hits in both directions including NR4A1, PDE1A, HOXB9, RB1, PCBP1 and LRRC10B are labeled. **e**, Long-range A-A contact frequencies for non-targeting control and selected hits. **f**, Log2 fold change of long-range A-B contact frequency versus -log10 FDR for each perturbation. Top three hits in both directions, including RFESD, HOXB9, FAM69B, C2CD2, CHD7 and FAM13C, are labeled. **g**, Long-range A-B contact frequencies for non-targeting control and selected hits. **h**, Log2 fold change of long-range B-B contact frequency versus -log10 FDR for each perturbation. Top hits in both directions, including FOS, NR4A1, DDX24 and MYBPH, are labeled. **i**, Long-range B-B contact frequencies for non-targeting control and selected hits. **j**, Log2 fold change of overall inter-TAD distances versus -log10 FDR for each perturbation. Top three hits in both directions, including PCBP1, RB1, CHD7, GLDC, HOXB9 and CUL1, are labeled. **k**, Overall inter-TAD distances for non-targeting control and selected hits. **l**, Log2 fold change of individual overall inter-TAD distances in chr22 for selected hits. P values in **c** and **k** were calculated by two-sided Wilcoxon signed rank test. P values in **e**, **g** and **i** were calculated by two-sided Wilcoxon rank sum test. In all box plots throughout the manuscript, the boxes cover the 25^th^ to 75^th^ percentiles, the whiskers cover the 10^th^ to 90^th^ percentiles, and the line in the middle of the boxes represents the median value. For all relevant panels, significance is represented as *p<0.1. **p<0.05. ***p<0.01.

We next explored the regulation of chromatin folding conformations at the length scale of A-B compartmentalization by measuring long-range chromatin contact frequencies for each perturbation within the same cells screened above. We defined two TADs spaced less than 500 nm apart as in contact with each other, and derived the long-range contact frequency between non-adjacent TADs along the chr22 genomic map. As A-B compartment organization stems from long-range chromatin contact^36^, we further categorized the long-range contacts as contacts between compartment A regions (A-A), contacts between compartment B regions (B-B), and inter-compartmental contacts between A and B regions (A-B). We observed that loss of NR4A1, PDE1A or the homeobox transcription factor HOXB9 increased the long-range A-A contact frequency, while knocking out RB1, PCBP1 or LRRC10B showed the opposite phenotype (Fig. 2d,e). Knocking out RFESD, HOXB9 or FAM69B increased the frequency of inter-compartmental A-B contacts, while knockout of C2CD2, the chromatin remodeler CHD7 or FAM13C decreased A-B contacts (Fig. 2f,g). Finally, knocking out the AP-1 transcription factor subunit FOS, NR4A1, or the helicase DDX24 increased the long-range B-B contact frequency, while knocking out MYBPH decreased B-B contact frequency (Fig. 2h,i).

Finally, we integrated all pairs of inter-TAD distances to measure and compare the overall compactness of the chr22 chromosome territory. The average fold changes of all 351 inter-TAD distances among the 27 TADs were calculated. We found that knocking out PCBP1, RB1, or CHD7 decompacted chr22 (Fig. 2j-l), while GLDC, HOXB9 or C19orf57 knockout resulted in chr22 compaction (Fig. 2j,k). Intriguingly, RB1 is known to promote the formation of senescence-associated heterochromatin foci^60^, a highly compacted whole-chromosome conformation. Our results show that RB1 knockout decompacts chr22 across multiple scales, including increased adjacent TAD distance (Fig. 2a-c), reduced long-range A-A contact frequency (Fig. 2d,e), and overall decompacted chromosome territory (Fig. 2j-l). The high-content readouts of our screen are therefore in agreement with this prior association but, importantly, provide new insights into the scales at which RB1 and the other hits impact chromatin organization.

We noticed that several chromatin folding regulators identified above, such as RB1, NR4A1, GLDC, HOXB9, PCBP1 and CHD7, were called as top hits in more than one architectural category. This observation led us to ask if chromatin folding regulators in general tend to affect multiple chromatin architectures across different length scales. To this end, we quantified the regulatory effects of each top hit on all five architectural features analyzed above (adjacent TAD distances, A-A, B-B and A-B interactions, and whole chromosome compaction). The results showed that most top hits significantly affect chromatin folding in more than one architectural category (Fig. 3a). In general, the top hits can be classified into chromatin compactors that reduce inter-loci distances and increase contact frequencies, and chromatin decompactors with the opposite function, although the extents of the regulatory effects often differ between categories for a given regulator (Fig. 3a). These observations indicate 3D genome regulators often have a multi-scale effect, but may also preferentially control architectures at certain scales.

**Fig. 3.**
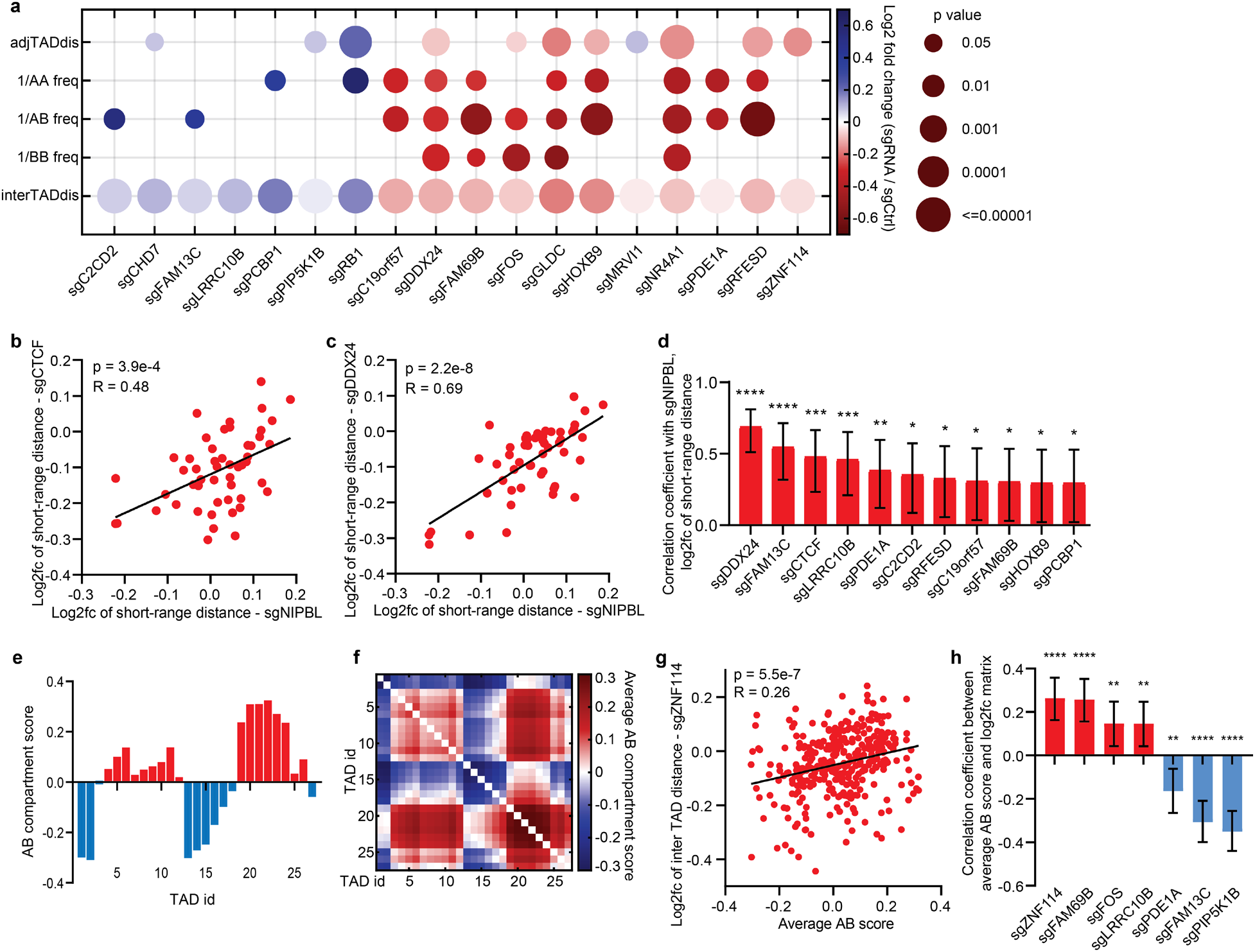
Characterization of the regulators of multi-scale chromatin folding. **a,** Fold change (bubble color) and significance (circle size) of multi-scale chromatin folding phenotypes of top hits. Phenotypic changes with p < 0.05 are not shown. **b**, Correlation of log2 fold change (log2fc) of short-range inter-TAD distance (defined as spatial distances between genomic regions that are less than 3Mb apart) between sgNIPBL (x axis) and sgCTCF (y axis). **c**, Correlation of log2fc of short-range inter-TAD distance between sgNIPBL (x axis) and a representative top hit sgDDX24 (y axis). **d**, Top hits significantly correlated with NIPBL in log2fc of short-range inter-TAD distance upon knockout. **e**, A-B compartment score profile of Chr22. **f**, Matrix of average A-B compartment scores of pairs of TADs. **g**, Correlation between the log2fc of inter-TAD distance upon ZNF114 knockout and the average A-B compartment score of the TADs. **h**, Top hits with 3D genome effects (log2fc of inter-TAD distance upon knockout) significantly correlated with the AB compartment score matrix. Error bars in **d** and **h** represent 95% confidence intervals. Stars represent the significance of the correlation: *p<0.05, ** p<0.01, ***p<0.001, ****p<0.0001.

### Correlation analyses link new regulators to known 3D genome mechanisms

The high content nature of our screen offers whole distance matrices for individual perturbations, allowing for correlation analyses between the 3D genome regulatory effects of the new regulators and those of previously identified mechanisms. Exploiting this capacity, we first quantified the correlations between each new regulator and NIPBL in controlling short-range (< 3 Mb) chromatin distances to detect potential mechanistic associations with the loop extrusion mechanism^61^. As a positive control, CTCF-knockout-induced short-range distance changes significantly correlated with changes upon NIPBL knockout (Fig. 3b). Among the 18 top hits, 10 showed significant correlations with NIPBL (Fig. 3c,d), suggesting at least partial interactions with the loop extrusion mechanism.

We next asked whether the regulatory effects of the top hits are associated with or mediated by the A-B compartmentalization scheme. To this end, we converted the one-dimensional AB compartment score profile to a 2D matrix by calculating the average AB compartment score between pairs of TADs (Fig. 3e,f), and measured the correlations between this average AB score matrix and the log2 fold change matrices of inter-TAD distances for individual top hits. 7 out of the 18 top hits showed significant correlations in this analysis, among which ZNF114, FAM69B, FOS and LRRC10B knockout led to distance changes positively correlated with the average AB score, whereas PDE1A, FAM13C and PIP5K1B knockout showed distance changes negatively correlated with the average AB score (Fig. 3g,h). These results indicate that a subset of our identified top hits at least partially interact with or are modulated by the A-B compartmentalization mechanism.

### CHD7 is a long-range chromatin compactor

As we examined our hits for each chromatin organization phenotype, CHD7 drew our attention because it regulated chromatin conformation on different scales in opposing ways. CHD7, or chromodomain helicase DNA binding protein 7, is a chromatin remodeler known to promote local chromatin openness and is associated with CHARGE syndrome^25–27^. The results of our screen showed that CHD7 knockout significantly reduced long-range A-B contact frequencies (Fig. 2f,g) and resulted in decompaction of the whole chr22 territory (Fig. 2j,k). To validate this large-scale chromatin-compaction function of CHD7 using an orthogonal method, we performed siRNA knockdown of CHD7 in the same A549-Cas9 cell line. Western blot analysis confirmed that the CHD7 protein level was reduced by >90% (Extended Data Fig. 3a). Chromatin tracing of chr22 in the CHD7 knockdown (siCHD7) and control (siCtrl) cells showed that while the compartment identities of TADs were largely identical between siCtrl and siCHD7 (Extended Data Fig. 3b,c), and the A-B compartments were spatially positioned in a conventional, polarized manner^50^ in both cases (Extended Data Fig. 3d), A-A, B-B, and A-B contacts were less frequent in siCHD7 compared to siCtrl cells (Extended Data Fig. 3e). CHD7 knockdown also caused overall chromosome territory decompaction, represented by a global increase of inter-TAD distances (Fig. 4a and Extended Data Fig. 3f) and larger radii of gyration of the whole chromosome traces (Extended Data Fig. 3g). The spatial regulatory effect of CHD7 appeared stronger at long range (defined as spatial distances between genomic regions that are more than 3Mb apart) compared to short range (defined as spatial distances between genomic regions that are less than 3Mb apart) (Fig. 4b). Altogether, these results agree with the phenotypes we observed for CHD7 in our knockout screen, confirming that this factor facilitates chromatin compaction especially at long range.

**Fig. 4.**
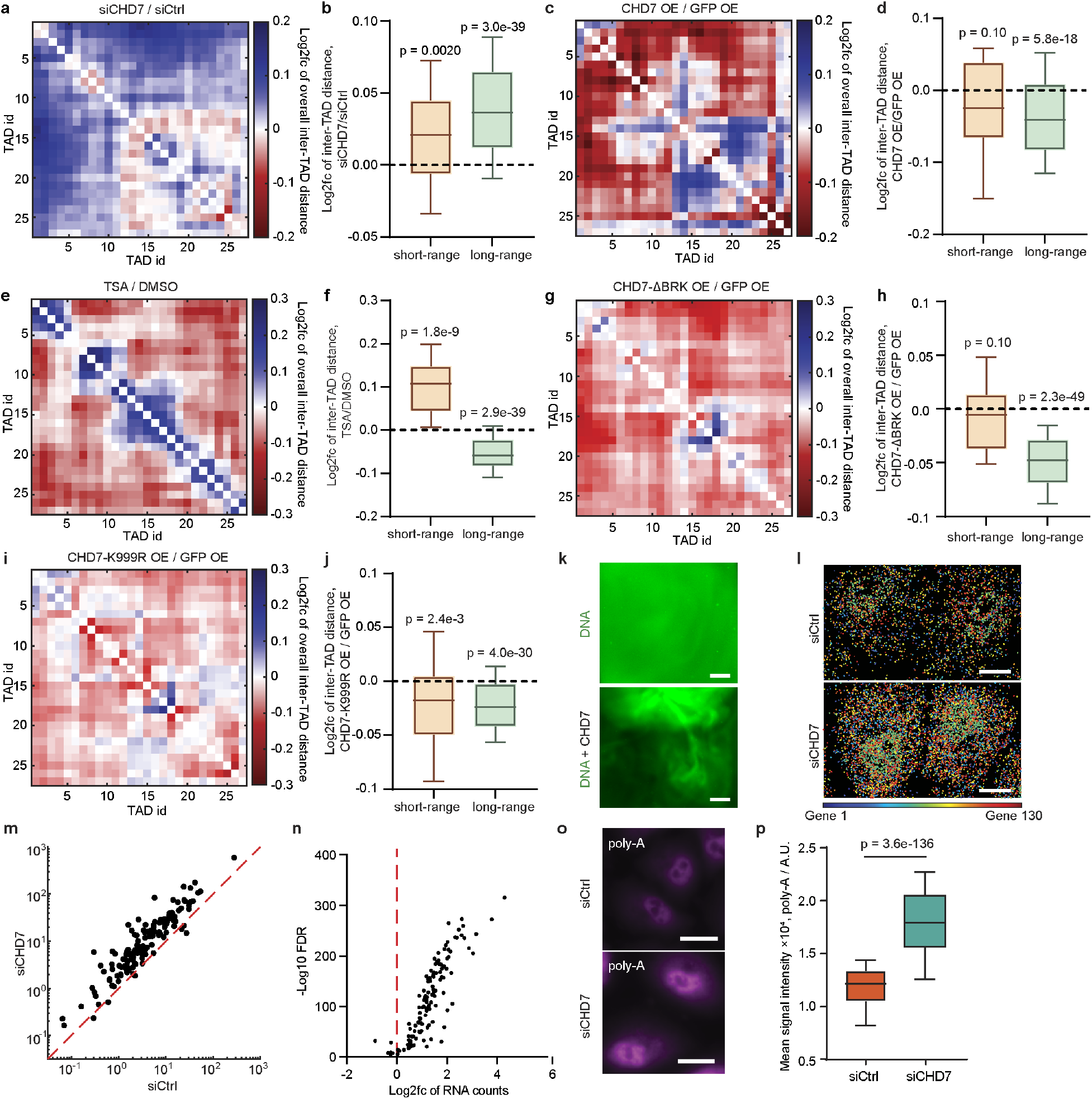
CHD7 is a long-range chromatin compactor that globally suppresses gene expression. **a**, Log2 fold change of inter-TAD distance of siCHD7 compared to siCtrl. Number of traces analyzed: 3,558 (siCtrl) and 4,134 (siCHD7). **b**, Log2 fold change of short-range (defined as spatial distances between genomic regions that are less than 3Mb apart) and long-range (defined as spatial distances between genomic regions that are more than 3Mb apart) inter-TAD distances between siCHD7 and siCtrl. **c**, Log2 fold change of inter-TAD distance of CHD7 overexpression compared to GFP overexpression. Number of traces analyzed: 3,157 (GFP OE) and 1,174 (CHD7 OE). **d**, Log2 fold change of short-range and long-range inter-TAD distances between CHD7 and GFP overexpression. **e**, Log2 fold change of inter-TAD distance of TSA-treated cells compared to DMSO-treated cells. Number of traces analyzed: 1,214 (DMSO) and 2,223 (TSA). **f**, Log2 fold change of short-range and long-range inter-TAD distances between cells with TSA and DMSO treatment. **g**, Log2 fold change of inter-TAD distance of CHD7-ΔBRK (BRK domain deletion) overexpression compared to GFP overexpression. Number of traces analyzed: 3,415 (CHD7-ΔBRK OE) and 2,164 (GFP OE). **h**, Log2 fold change of short-range and long-range inter-TAD distances between CHD7-ΔBRK OE and GFP OE. **i**, Log2 fold change of inter-TAD distance of CHD7-K999R overexpression compared to GFP overexpression. Number of traces analyzed: 2,045 (CHD7-K999R) and 2,164 (GFP OE) **j**, Log2 fold change of short-range and long-range inter-TAD distances between CHD7-K999R OE and GFP OE. All chromatin tracing experiments in this figure were done in the A549 cell background, targeting chr22. P values were calculated by two-sided Wilcoxon signed rank test. **k**, Representative images of dye-labeled lambda DNA without/with purified CHD7. Scale bar: 500 μm. **l**, Spatial distribution of 130 genes decoded by RNA MERFISH in siCtrl and siCHD7 cells. Two representative cells are shown for each condition. Scale bar: 10 μm. **m**, Average RNA counts per cell for each gene in siCHD7 versus siCtrl cells. The red dashed line represents the x=y line. **n**, -Log10 false discovery rate (FDR) versus log2 fold change (log2fc) of average RNA counts per cell for each gene from siCtrl to siCHD7. Number of cells analyzed: 1,979 (siCtrl) and 1,186 (siCHD7) in **m** and **n**. **o**, Representative cell images of poly-A stain for siCtrl and siCHD7 cells. Scale bar: 20 μm. **p**, Mean fluorescent intensity of poly-A stain in individual nuclei of siCtrl and siCHD7 cells. P value was calculated by two-sided Wilcoxon rank sum test. Number of nuclei analyzed: 666 (siCtrl) and 594 (siCHD7).

To further validate the regulatory effect of CHD7 on chromatin organization, we overexpressed CHD7 in the A549-Cas9 cell line. In comparison to control GFP overexpression using the same vector, CHD7 overexpression significantly compacted chromatin and promoted chromatin contacts across larger length scales: While the A-B compartment identities of TADs and the polarized arrangement of A-B compartments largely remained unchanged (Extended Data Fig. 4a-c), CHD7 overexpression led to significantly higher contact frequencies between A-A, B-B, and A-B compartment regions (Extended Data Fig. 4d), and significantly decreased global inter-TAD distances and radii of gyration (Fig. 4c and Extended Data Fig. 4e,f), indicating overall chromatin compaction. Similar to the CHD7 knockdown scenario, the compaction effect upon CHD7 overexpression is more significant at long range (Fig. 4d). Together, these data further validated the findings from our image-based pooled screen, and show that CHD7 specifically promote long-range chromatin compaction and contact.

To investigate if the chromatin compaction function of CHD7 applies to a different cell background and different genomic context, we performed siRNA knockdown of CHD7 in human RPE-1 cells and conducted chromatin tracing on chromosome 21 (chr21). CHD7 knockdown caused an overall increase of inter-TAD distance (Extended Data Fig. 5a,b) and an increase of the radius of gyration of chr21 territory (Extended Data Fig. 5c), which were mainly contributed by long-range chromatin decompaction (Extended Data Fig. 5d). CHD7 knockdown also led to decreased contact frequency between A-A, A-B and B-B interactions (Extended Data Fig. 5e). As CHARGE syndrome is known as a neural crest disease and the multiple organs affected by the syndrome are derived from neural crest progenitor cells in early development^24^, we further tested whether siCHD7 affects long range chromatin compaction in neural crest progenitor cells differentiated from cultured human embryonic stem cells. Western blots confirmed the neural crest cell identity and effective knockdown of CHD7 (Extended Data Fig. 6a). Chromatin tracing analyses confirmed that siCHD7 in neural crest cells caused long range chromatin decompaction (Extended Data Fig. 6b,c). Together, these results indicated that the chromatin compaction function of CHD7 exists in distinct genomic contexts and cell lines, including the cell type of origin of CHARGE syndrome.

### CHD7 directly compacts chromatin at long-range independently of its chromatin remodeling activity and globally represses gene expression

CHD7 is known to promote local chromatin accessibility and regulate transcription^27^ – while it is perhaps not surprising that CHD7 has an effect on genome organization in general, it was unintuitive and unexpected to discover that it promotes *compaction* of higher-order chromatin architectures. CHD7 belongs to the chromodomain helicase DNA-binding protein family, which are ATP-dependent chromatin remodelers^62^. It contains two chromodomains, an ATPase domain, and a SLIDE-SANT domain that are important for nucleosome sliding, and two BRK domains close to C terminus of largely unknown function^26^. Consistent with previous reports in other cell contexts^63^, we confirmed that CHD7 binds to a broad range of targets as shown by CUT&RUN (Extended Data Fig. 7a-c). Intriguingly, a recent report showed that in the context of DNA damage, CHD7 binds to the damaged site and initially decompacts chromatin, and then recruits histone deacetylases (HDAC) to re-compact chromatin through histone deacetylation^64^. To investigate the possibility that CHD7 could compact chromatin through regulating HDAC activity in the context of our undamaged cells, we treated wild-type cells with an HDAC inhibitor Trichostatin A (TSA). Immunofluorescence confirmed that TSA treatment increased global histone acetylation (Extended Data Fig. 8a), and chromatin tracing of chr22 showed that HDAC inhibition led to chromatin decompaction at short scales, including between adjacent TADs (Extended Data Fig. 8b-d), but had the opposite effect at long range (Fig. 4e,f and Extended Data Fig. 8e-g). Thus, the long-range chromatin compaction activity of CHD7 cannot be explained by its known HDAC recruitment capability.

It has been shown that CHD7 can directly bind CTCF through the BRK domains^65^. Indeed, our CUT&RUN profiling of CHD7 and CTCF showed that CHD7 binding peaks across the genome overlap with a subset of CTCF binding peaks in the A549-Cas9 cell line (Extended Data Fig. 7d). A previous report showed that CTCF and cohesin binding peaks along the genome and loop extrusion stripe features in Hi-C map were not affected by CHD7 knockout^27^, suggesting that CHD7 does not modulate CTCF binding to the genome or loop extrusion. To test if the chromatin compaction effect of CHD7 depends on its interaction with CTCF, we performed chromatin tracing on cells overexpressing a mutant CHD7 lacking the BRK domains, and compared the traces to those from control GFP overexpressing cells. Chromatin tracing showed that the BRK mutant CHD7 overexpression still led to decreased inter-TAD distances at long range (Fig. 4g,h). These results indicate that the long-range chromatin compaction function of CHD7 does not rely on its interaction with CTCF.

We then asked if the ATP-dependent chromatin remodeling activity of CHD7 is needed for its long-range chromatin compaction function by overexpressing a previously established mutant CHD7 with the lysine residue at position 999 in the ATP-binding motif mutated to arginine (K999R), which lacks the chromatin remodeling activity^26^. Chromatin tracing results in comparison to control GFP overexpression showed that CHD7 with the K999R mutation is still capable of long-range chromatin compaction (Fig. 4i, j), indicating that the ATP-dependent chromatin remodeling activity is dispensable for the new chromatin compaction function. In other words, the observed long-range chromatin compaction is not a secondary effect of CHD7 chromatin remodeling. Finally, to check whether CHD7 is self-sufficient in compacting/condensing DNA in an ATP-independent manner, we purified CHD7 and tested its ability to condense DNA *in vitro*. The results showed that dye-labeled lambda DNA can be effectively condensed with purified CHD7 without ATP or other proteins, forming large scale DNA condensates (Fig. 4k). Together, our data indicate that CHD7 directly compacts DNA independently of chromatin remodeling.

We hypothesized that the large scale chromatin compaction function of CHD7 may globally regulate gene expression. To test this hypothesis, we first performed bulk RNA sequencing using siCHD7 and control A549-Cas9 cells. Consistent with previous reports in other cell contexts^63^, our bulk RNA-seq analyses showed that CHD7 knockdown resulted in diverse gene expression changes (up- and down-regulation) in many cellular programs (Extended Data Fig. 7e,f). However, we reasoned that the read normalization step in routine bulk RNA-seq analysis pipeline would eliminate any global expression differences between samples. To detect potential systematic differences in gene expression levels between siCHD7 and control cells, we performed RNA multiplexed error-robust FISH (MERFISH) targeting 130 randomly selected genes using a previously reported probe library^66^, and measured the RNA copy numbers per cell of the target genes. Based on the MERFISH data, 124 out of the 130 genes showed significant up-regulation upon CHD7 knockdown (Fig. 4l-n). To sample transcripts from more genes, we further performed fluorescence labeling of all poly-A transcripts in siCHD7 and control cells using poly-dT probes, and quantified the fluorescence intensities in the cell nuclei to better estimate nascent transcription levels. siCHD7 cells exhibited significantly higher fluorescence intensities than control cells (Fig. 4o,p). These results indicate that CHD7 globally suppresses gene expression and helps establish lower baseline expression levels for genes.

### Identification of regulators of chromosomal radial positioning and morphological properties of the nucleus

Chromatin association with the nuclear lamina is functionally linked to gene silencing^67, 68^. Gene-rich chromosomes (such as chr22) are known to be preferentially located towards the nuclear interior^69^. Localization to the interior of the nucleus in interphase can also protect chromosomes from segregation errors during mitosis^70^. To screen for factors that regulate chromatin-lamina association, we incorporated DAPI staining of whole nuclei together with our multi-scale chromatin tracing described above in the context of our CRISPR screen. We used the edges of the whole nuclear staining patterns to approximate lamina positions, and identified gene knockout hits that affected the distance between chr22 and the nuclear lamina (Fig. 5a-c). We found that knocking out CUL1, PRSS22, or MAB21L2 brought chr22 closer to the nuclear lamina, as shown by shorter distances both between the centroid of the chromosome to the lamina (Fig. 5a,b) and between each individual TAD to the lamina (Fig. 5c), while LRRC10B, DDX21, or HMGA2 knockout moved chr22 towards the nuclear interior (Fig. 5a-c).

**Fig. 5.**
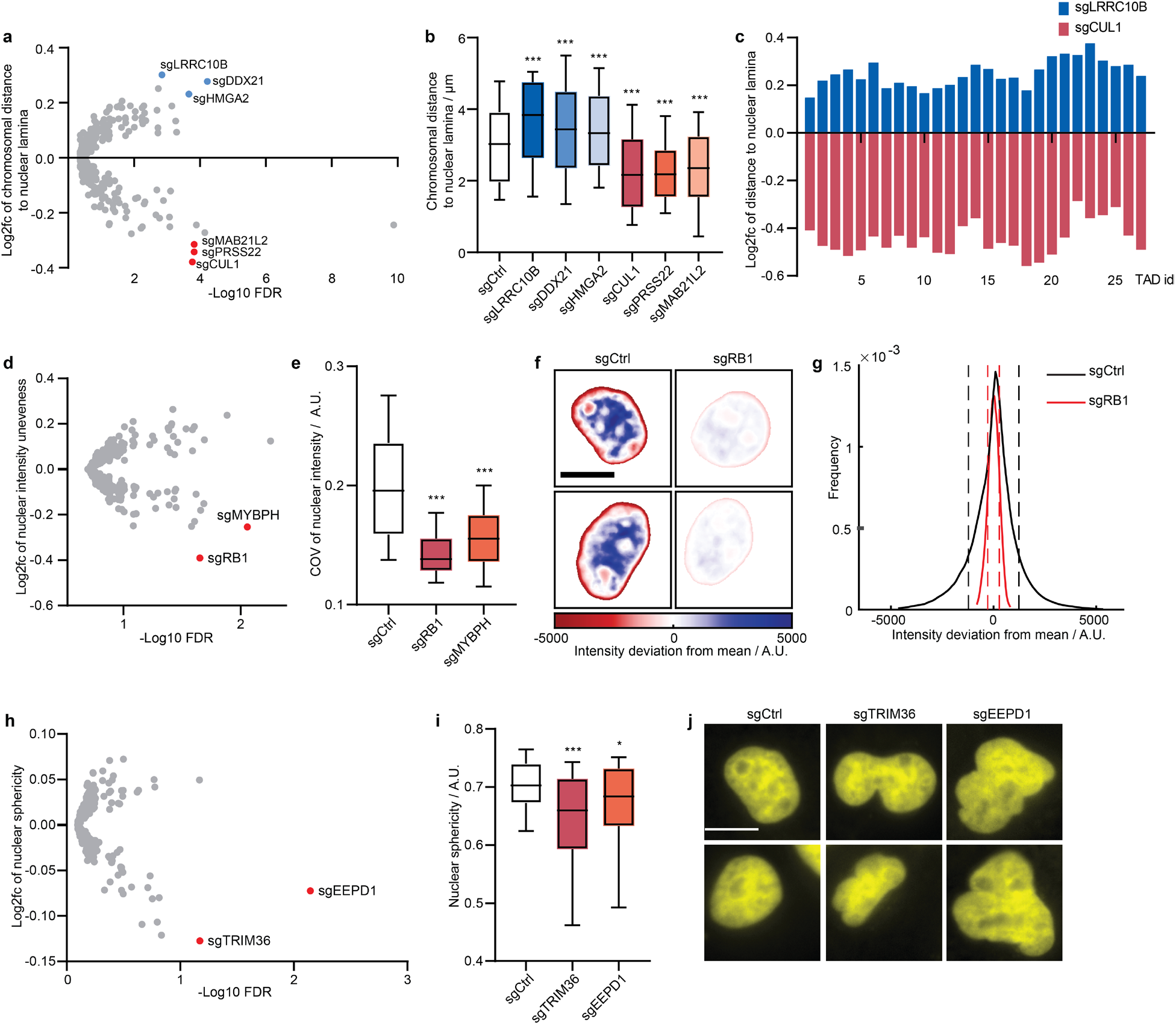
Perturb-tracing screen identified hits that regulate chromosome association with nuclear lamina and the morphological properties of the nucleus. **a**, Log2 fold change of chromosome distance to nuclear lamina versus -log10 FDR. Top hits in both directions, including LRRC10B, DDX21, HMGA2, CUL1, PRSS22 and MAB21L2, are labeled. **b**, Chromosome distance to nuclear lamina of non-targeting control and selected hits. **c**, Log2 fold change of distances between each TAD on chr22 to nuclear lamina of selected hits. **d**, Log2 fold change of nuclear intensity unevenness (measured as coefficient of variation of nuclear voxel intensities) versus -log10 FDR. Top hits RB1 and MYBPH are labeled. **e**, Nuclear intensity unevenness of non-targeting control and selected hits. **f**, Heatmap of nuclear intensity deviation from mean intensity of representative nuclei from non-targeting control (left column) and selected hit sgRB1 (right column). Scale bar: 10 μm. **g**, Voxel intensity distribution of all nuclei from non-targeting control (black curve) and selected hit sgRB1 (red curve). Dashed lines indicate the standard deviations of the indicated distributions. **h**, Log2 fold change of nuclear sphericity versus -log10 FDR. Top hits TRIM36 and EEPD1 are labeled. **i**, Nuclear sphericity of non-targeting control and selected hits. **j**, Representative nuclei images of non-targeting control (left column) and selected hits that regulate nuclear sphericity, TRIM36 (middle column) and EEPD1 (right column). Each column contains the DAPI staining of two representative cells from the indicated perturbation. Scale bar: 10 μm. P values in **b**, **e** and **i** were calculated by two-sided Wilcoxon rank sum test.

Even on its own, nuclear DAPI staining bears surprisingly rich information regarding the nuclear organization that can be used to distinguish cell states^71, 72^. Chromatin is distributed unevenly in the cell nucleus, and the “texture” of nuclear staining pattern is often used as a diagnostic marker of cancer^73^. Moreover, cells with abnormal, non-spherically shaped nuclei are often seen in cancer and aging and may indicate genome instability^74–76^. By analyzing changes in DAPI staining caused by each perturbation in our screen, we found that knockout of RB1 or MYBPH reduced the unevenness of nuclear DAPI staining, generating patterns with more homogeneous intensity within each nucleus (Fig. 5d-g), whereas a more heterogenous intensity pattern with chromatin condensates is often associated with cancer^73^. In addition, knocking out TRIM36 and EEPD1 decreased the sphericity of the nuclei and led to multi-lobed nuclei shapes (Fig. 5h-j).

### A general link between chromatin compaction regulation and nuclear sphericity

To test if the nuclear phenotypes are fully orthogonal or at least partially linked to the chromatin folding phenotypes analyzed before, we calculated the correlations among the changes of different chromatin folding and nuclear organization features upon perturbations of the top hits. As expected, all chromatin folding features (adjacent TAD distances, A-A, B-B, and A-B contact frequencies, and inter-TAD distances) are significantly correlated with each other (Fig. 6a), consistent with the common muti-scale and multi-faceted effects of the chromatin folding regulators (Fig. 3a). Surprisingly, nuclear sphericity also showed significant correlations with chromatin folding features across length scales and categories (Fig. 6a). In general, chromatin compaction (decreased inter-loci distance or increased contact frequency) is associated with a more spherical nuclear shape (Fig. 6a). To validate this observation, we measured and compared the nuclear sphericities of siCHD7 and control A549-Cas9 cells. Indeed, CHD7 knockdown led to a less spherical and more multi-lobed nuclear shape (Fig. 6b,c). To interpret this result, we performed simulation of a minimal chromatin polymer model. We showed that reduced monomer-monomer interaction strength (less chromatin compaction/interaction) can lead to a less globular chromatin folding organization, and a bounding envelope surrounding the polymer will in turn adopt a more multi-lobed shape with lower sphericity (Fig. 6d). Overall, our experimental results suggest a general link between chromatin compaction and nuclear shape, and our simulation supports a model that chromatin compaction mediates nuclear morphology.

**Fig. 6.**
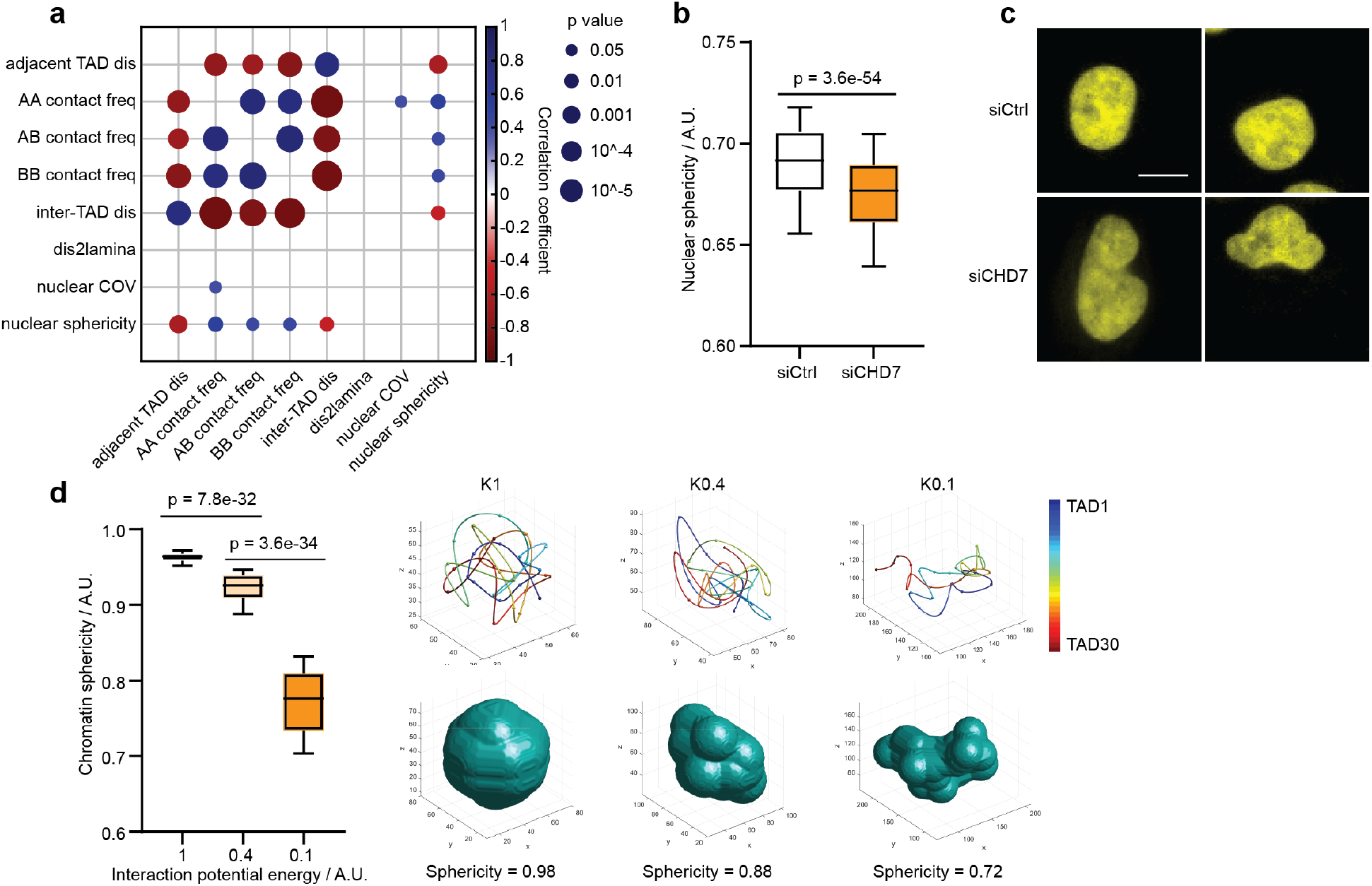
A link between chromatin compaction and nuclear shape. **a,** Correlation coefficients (bubble color) and significance of correlations (bubble size) between pairs of 3D genome/nucleome features calculated using all top hits. **b**, Nuclear sphericities of siCtrl and siCHD7 A549-Cas9 Cells. Number of cells analyzed: 1,156 (siCtrl) and 1,412 (siCHD7). **c**, Representative DAPI images of siCtrl and siCHD7 A549-Cas9 cells. **d**, Simulated chromatin polymer folding conformations and the corresponding bounding envelop sphericities at different chromatin self-interaction energies (K = 1, 0.4 or 0.1). Lower energy corresponds to weaker chromatin interaction. N = 100 simulated conformations for each energy. P values in **b** and **d** were calculated by two-sided Wilcoxon rank sum test.

In summary, our image-based Perturb-tracing screen using pooled CRISPR perturbations, BARC-FISH and multi-scale chromatin profiling allowed us to systematically and simultaneously profile the effect of hundreds of candidate genes on many aspects of spatial genome organization, from short- and long-range chromatin folding and chromatin association with a nuclear landmark to whole-nucleus morphological characteristics. Overall, we identified 26 top candidate regulators of chromatin/nuclear organization from our screen of 137 selected genes (Supplementary Table 2), and present a generalizable platform that can be used to screen thousands of genes of interest in different cell types and conditions in future work.

## Discussion

In this work, we report the development of a high-throughput, high-content, image-based genetic screening platform termed Perturb-tracing and demonstrate its ability to systematically identify regulators of 3D genome folding architectures from short-range to long-range and global nuclear organization. The Perturb-tracing platform integrates a pooled CRISPR knockout screen comprising hundreds of candidate genes with chromatin tracing and the development of a cellular barcoding and *in situ* decoding technique termed BARC-FISH. Other FISH-based *in situ* genotyping techniques have previously been reported in genetic screens but are limited to bacterial applications and/or not integrated with 3D genomics^49, 77, 78^. Our BARC-FISH technique combines *in situ* amplification and robust readout of a short RNA barcode associated with a CRISPR gRNA in each cell, thereby allowing us to pair the genetic perturbation through CRISPR (cause) with the 3D genome alteration (effect) observed in each cell. The BARC-FISH barcode design has the capacity to simultaneously assess libraries of up to ∼5000 sgRNAs in pooled format with the current experimental design and can be further extended in the future with longer barcodes. Both BARC-FISH and chromatin tracing are based on highly multiplexed FISH, and thus are readily compatible with each other on the same sample. Importantly, the phenotypic readout of this screening platform is information-rich, allowing for categorization of a gene of interest based on its effects on many different length scales and aspects of chromatin organization. Overall, our current work included ∼13,000 imaging target-perturbation combinations (420 gRNAs multiply 30 phenotypic imaging targets including 27 TADs, DAPI, total protein, and Geminin stains). The rich information and high-content nature of the screen also readily enables detection of associations between new regulators and known 3D genome organizers, and discovery of potentially functional linkages and co-regulation mechanisms between different 3D genome/nucleome features. We expect the entire screen methodology to be broadly adaptable to many other cell types and biological contexts.

In our initial screen of 137 candidate genes with multiple sgRNAs per gene, we identified 26 previously unknown or incompletely known regulators of 3D genome and nuclear organization features across multiple length scales, including the architecture of adjacent TADs, A-B compartment organization, entire chromosome conformation and localization, and whole nucleus morphology. We observed and validated a long-range chromatin compaction function of CHD7, which was striking because of its incongruence with the opposite chromatin accessibility-promoting effect of this protein at short scale. This observation suggests that CHD7-targeted genomic regions may be subject to a new type of “structural poising”, containing features of both open chromatin and compact chromatin at different length scales. We further showed that the long-range chromatin compaction function of CHD7 is a direct effect independent of its regulation of HDAC activity, its interaction with CTCF, and its chromatin remodeling function, indicating distinct mechanisms governing short- and long-range activities of CHD7. Our identification and characterization of CHD7 in this paper demonstrates how our screening methodology can generate new hypotheses with sufficient information to guide targeted follow-up experiments. We anticipate that the remaining hits from our screen, as well as hits from future broader screens encompassing a wider range of candidate genes, will provide a rich resource for discovery of new chromatin regulators at multiple scales and their mechanisms of action.

Interestingly, a congenital disorder known as CHARGE syndrome (named for its associated phenotypes of coloboma, heart defects, atresia, growth retardation, genital abnormalities, and ear abnormalities^25^) is highly attributed to mutations in CHD7^79^, and the new functional insights we gained through this work could advance understanding of how CHD7 contributes to these developmental abnormalities. Intriguingly, our results showed the CHD7 globally suppresses gene expression and contributes to a more spherical nuclear shape, both of which may be attributed to its long-range chromatin compaction function. A correct baseline of global gene expression and nuclear morphology may be particularly important in early development, when genes and cells are highly dynamic, and defects may lead to the many symptoms in complex diseases such as CHARGE syndrome. A close paralog of CHD7 known as CHD8 is implicated in autism spectrum disorder^80^, and while CHD8 was not represented in our screen, some of our insights and proposed mechanism for CHD7 function may translate to CHD8 as well.

The candidate 3D genome regulators identified in this work have diverse expression profiles in different tissues^81^ (Extended Data Fig. 9), and 24 out of 26 of our reported top hit genes are associated with disease states or human phenotypes (Supplementary Table 2). Our screen provides a potential mechanistic link between 3D genome regulation and the physiology/pathology of the diverse tissues and contexts. For example, NR4A1, an orphan nuclear receptor protein, has been found to be a signature of metastasis in certain primary solid tumors^82^. Our screen found it to be a top candidate in regulating chromatin compartment interactions (Fig. 2d,e and h,i). It is of great interest to investigate whether the chromatin conformation changes it mediates contribute to tumor metastasis. Our top candidates also included important genes in development, for example FOS, HOXB9 and MAB21L2. 3D genome reorganization has been shown to be crucial in regulating tissue-specific gene expression during development^83^. We are intrigued by how these genes may regulate development in part through 3D genome regulation. ZNF114 was shown to associate with MDC1 in a two-hybrid screen, a critical regulator of DNA double-strand break (DSB) repair^84^. Substantial data show that DSBs lead to a decompaction of heterochromatin and relocation to the periphery of heterochromatic domains^85^. Given its association with MDC1 and observed compaction upon knockout, ZNF114 may regulate the topological reorganization of the genome after genotoxic stress and the generation of DSBs. Besides the identification of novel 3D genome regulators, our work also raised new questions about previously known or speculated 3D genome regulators. For example, RB1 was known to recruit cohesin and condensin II complexes during DNA replication and mitosis for sister chromatid cohesion and chromosome condensation^86^. Our results showed a prominent role of RB1 in promoting higher-order chromatin compaction during G1, likely through its regulation of heterochromatin formation and maintenance^86^.

Overall, Perturb-tracing enables mapping of the chromatin organization “regulatome” at scale, which will deepen our understanding of the regulatory landscape of the genome and of the functions of genome architectures in diverse developmental and disease states.

---

**Extended Data Fig. 1.**
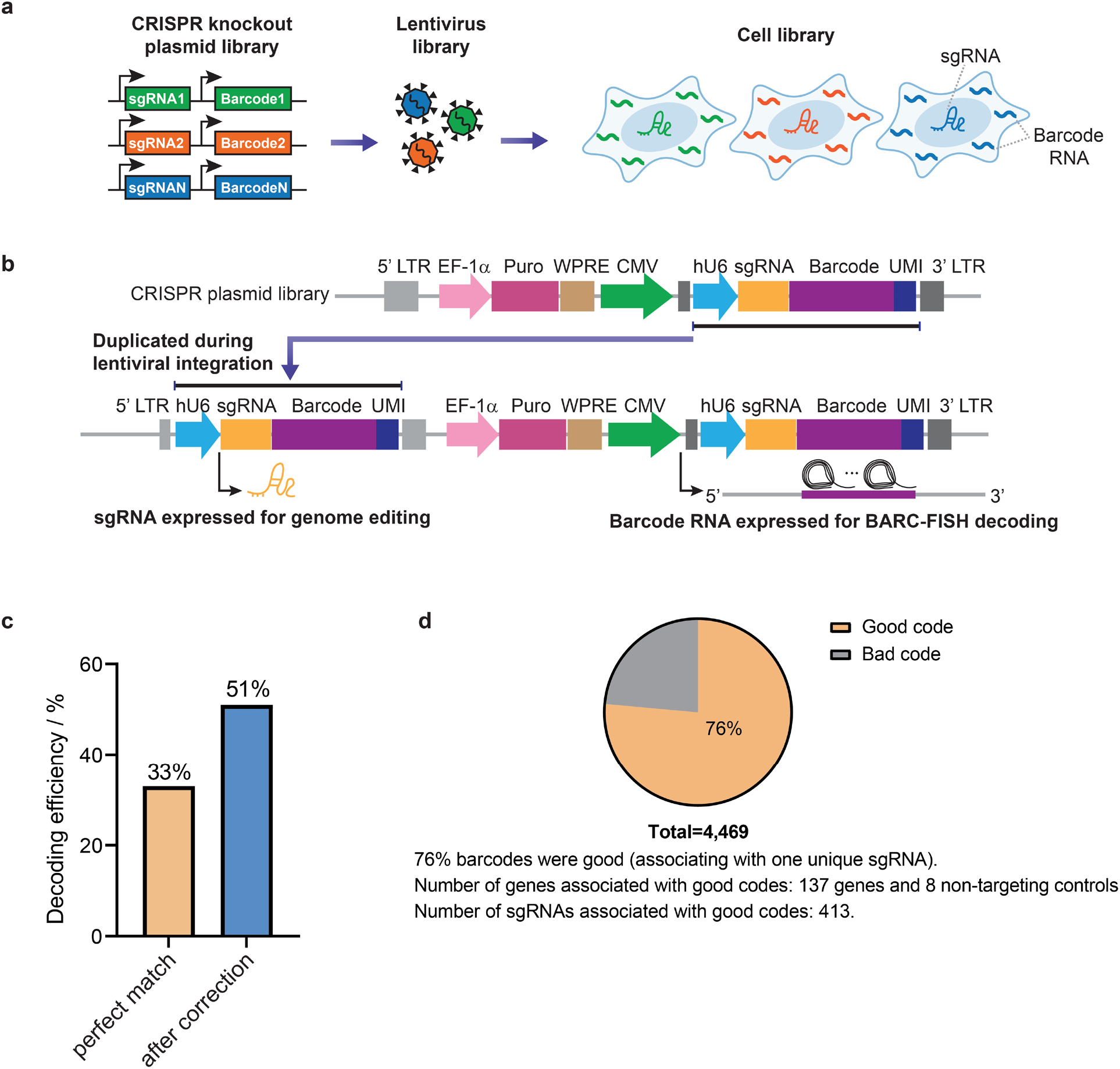
CRISPR screen library allows for sgRNA expression for genome editing and the barcode RNA expression for BARC-FISH decoding. **a**, Schematic of the construction of CRISPR screen library. A CRISPR knockout plasmid library containing sgRNA-barcode associations was constructed to generate a lentivirus library, which was transduced into human A549-Cas9 cells to produce a cell library. **b**, Design of the CRISPR screen plasmid and the lentiviral integration strategy. The sgRNA-barcode cassette was composed of human U6 promoter (hU6, blue), sgRNA (yellow), barcode (purple) and UMI (dark blue) sequences and placed within 3’ long terminal repeat (3’ LTR, dark gray), downstream of a strong RNA Pol II promoter (CMV, green). This cassette was duplicated and inserted within 5’ LTR (light gray) during lentiviral integration. Therefore, the cassette within 5’ LTR was able to express the sgRNA for genome editing, while the other copy was driven by CMV promoter to express a high level of barcode RNA for BARC-FISH decoding. Other elements on the plasmid including EF-1α promoter (pink), Puromycin resistance gene (magenta) and WPRE element (brown) were shown. **c**, BARC-FISH decoding efficiency in the CRISPR screen cell library. After BARC-FISH decoding procedure, the decoded barcodes were compared and matched to the barcodes determined by NGS. 33% of the imaged cells contained barcodes with perfect matches. After the error correction, 51% of the cells contained matched barcodes. **d**, Analysis of barcode quality determined by NGS. The sgRNA-barcode associations in the CRISPR screen cell library were determined by NGS (see Methods, “Detection of sgRNA-barcode associations in the cell library” section). In total, 4,469 barcodes were detected, among which 76% were good codes associating with one unique sgRNA. 413 sgRNAs targeting 137 genes and 8 non-targeting controls were found to be associated with these good codes. 412 of the 413 sgRNAs were observed in the image-based screen.

**Extended Data Fig. 2.**
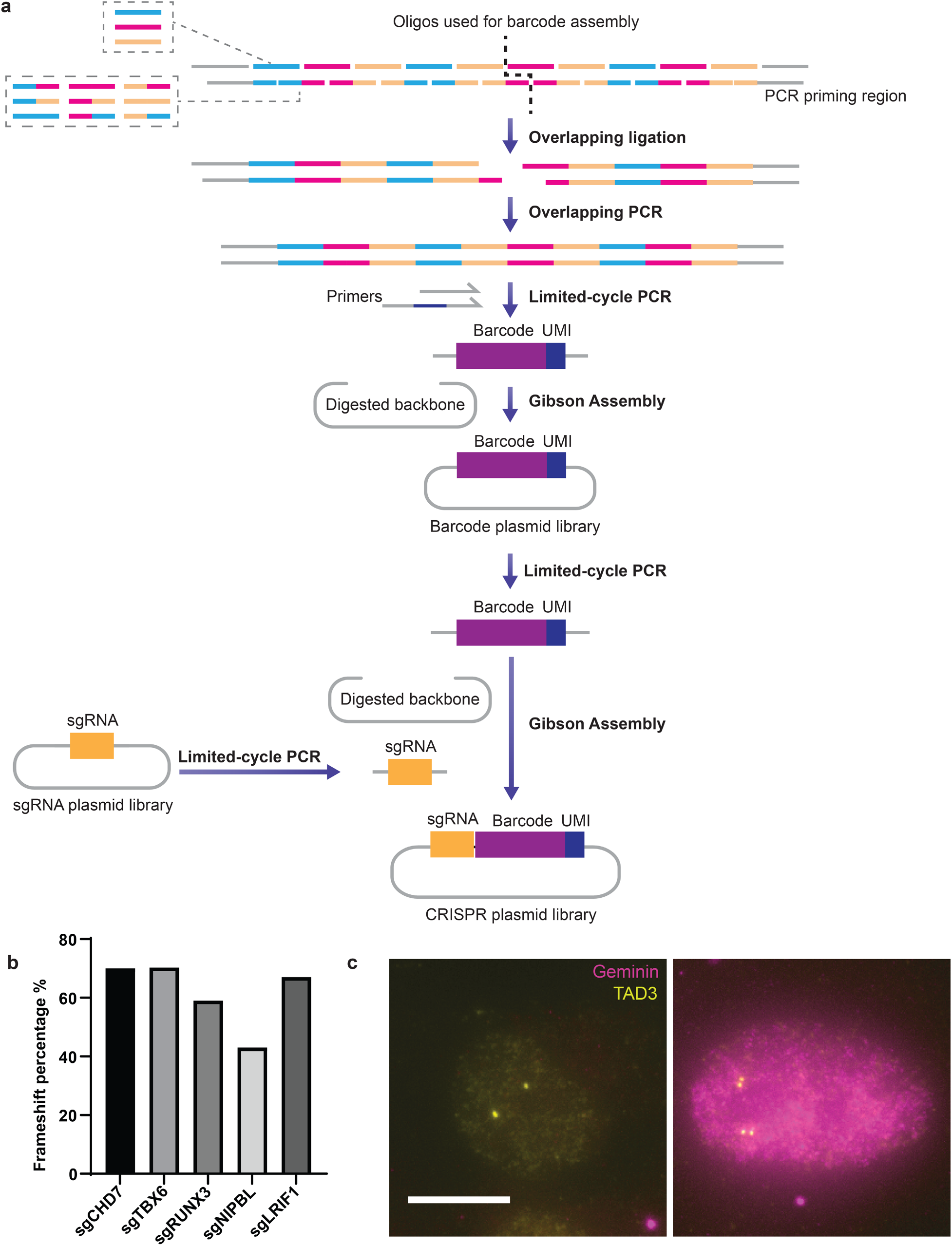
Cloning strategy of plasmid libraries, CRISPR knockout efficiency, and the Geminin-based cell cycle identification. **a**, The barcode plasmid library was assembled from individual oligos through overlapping ligation, overlapping PCR, limited-cycle PCR and Gibson Assembly (see Methods, “Barcode plasmid library construction” section). Each of the forward-strand oligos contained three alternative sequences (in the smaller gray dashed box), represented by three different colors cyan, magenta, and yellow. The overlapping oligos in the reverse strand contained 9 alternative sequences (in the larger gray dashed box). Oligos at the two ends carried PCR priming regions (straight gray lines). The barcode was divided into two halves which were subjected to overlapping ligation to form two double-stranded fragments. The two fragments were assembled by overlapping PCR to form a full-length barcode. The barcode was then amplified and added with UMI (unique molecular identifier) by limited-cycle PCR primers. The barcode-UMI fragments were inserted into a digested plasmid backbone through Gibson Assembly to construct the final barcode plasmid library. To clone the CRISPR screen plasmid library, sgRNA fragments and barcode-UMI fragments were amplified from the premade sgRNA plasmid library and barcode plasmid library respectively, through limited-cycle PCR. The sgRNA and barcode-UMI were then Gibson Assembled into a digested lentiviral plasmid backbone to generate the final CRISPR screen plasmid library (see Methods, “CRISPR screen plasmid library construction” section). The UMI was necessary for sequencing-based mapping of barcode-sgRNA associations (see Methods, “Next-generation sequencing (NGS) library preparation for mapping sgRNA-barcode associations”). **b**, Percentage of frameshift mutations of sgCHD7, sgTBX6, sgRUNX3, sgNIPBL and sgLRIF1. **c**, Two representative cells from the screen datasets were shown to demonstrate the Geminin staining strategy for G1 phase cell detection. Geminin antibody stain (magenta) is absent in a G1 phase cell (left), which showed two DNA FISH foci of TAD3 (yellow) of chr22. The S/G2 phase cell (right) is positive for Geminin stain and have four DNA FISH foci of TAD3 in two pairs, indicating replicated TAD3 DNA. Because Geminin and the yellow-green fiducial beads were imaged using the same laser channel, bead patterns were seen in both images (small, round magenta spots outside of the nuclei). Scale bar: 10 μm.

**Extended Data Fig. 3.**
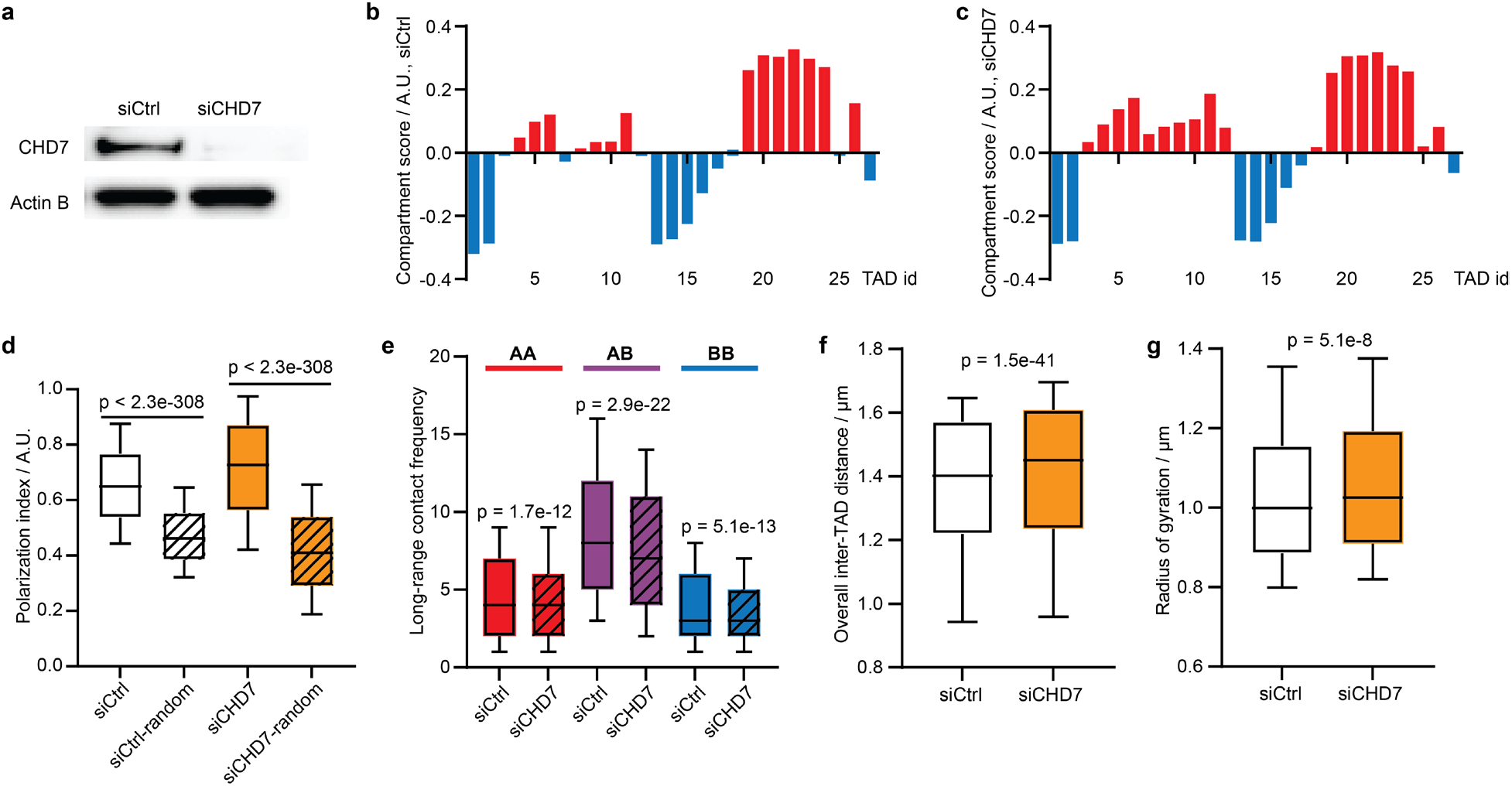
Validation of CHD7 perturbation phenotypes using RNA interference. **a**, Western blot of siCtrl- and siCHD7-treated A549-Cas9 nuclear extracts. Top: anti-CHD7 antibody; bottom: anti-Actin B antibody. **b**, A-B compartment profile of chr22 in siCtrl cells. **c**, A-B compartment profile of chr22 in siCHD7 cells. **d**, Polarization indices of chr22 A-B compartments of siCtrl (white) and siCHD7 (orange). Shadowed boxes show the polarization indices from randomized controls, where the compartment identities of TADs are scrambled. **e**, Compartmental contact frequencies of siCtrl and siCHD7 (shadowed) among A compartment regions (red), between A and B compartment regions (purple), and among B compartment regions (blue). **f**, Overall inter-TAD distance of siCtrl and siCHD7. **g**, Radii of gyration of siCtrl and siCHD7. P values in **d**, **e** and **g** were calculated by two-sided Wilcoxon rank sum test. P value in **f** was calculated by two-sided Wilcoxon signed rank test.

**Extended Data Fig. 4.**
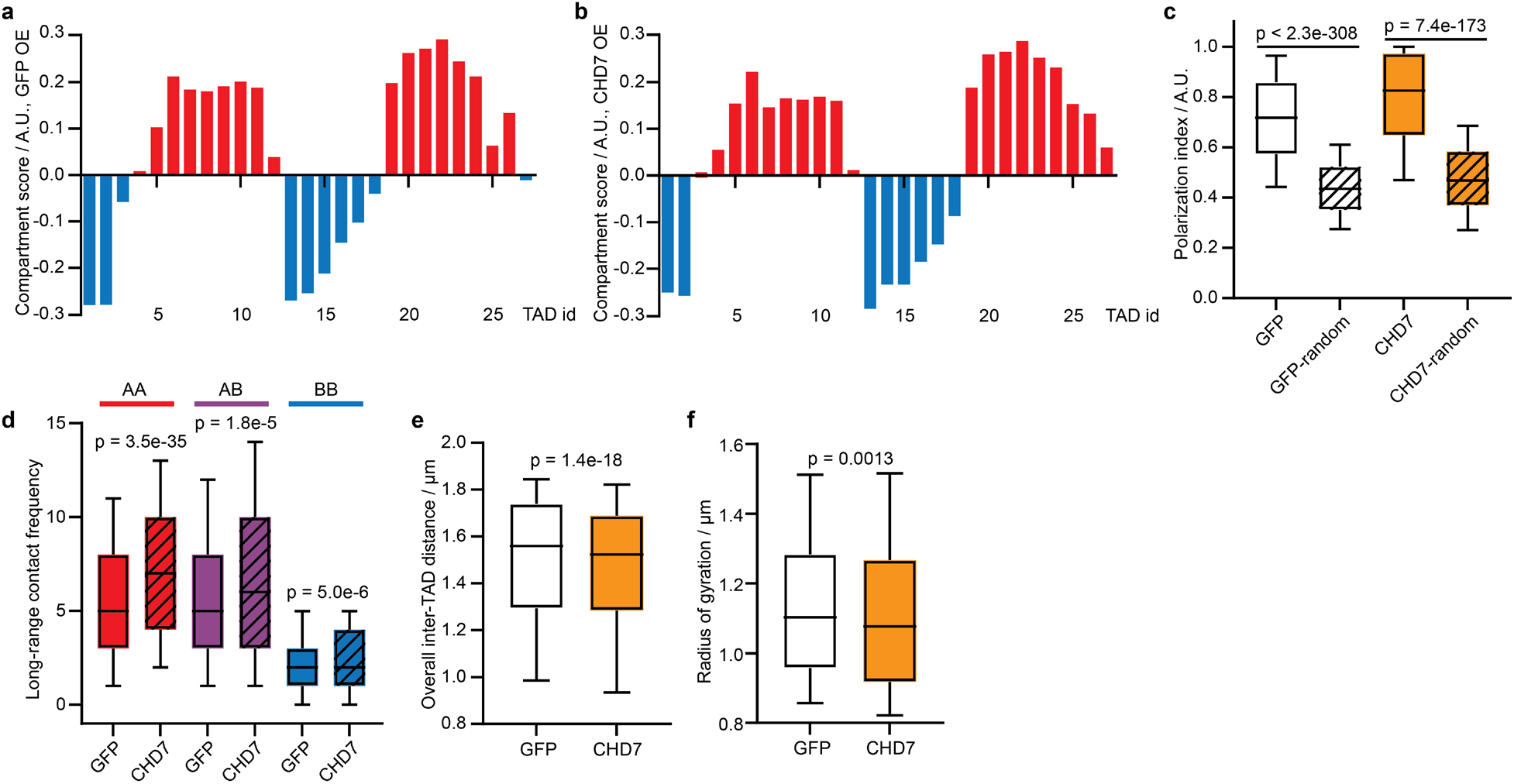
Validation of CHD7 perturbation phenotypes using overexpression. **a**, A-B compartment profile of chr22 in A549-Cas9 cells with GFP overexpression. **b**, A-B compartment profile of chr22 in A549-Cas9 cells with CHD7 overexpression. **c**, Polarization indices of cells with GFP (white) and CHD7 (orange) overexpression and the corresponding randomized controls (shadowed). **d**, Compartmental contact frequencies of cells with GFP of CHD7 (shadowed) overexpression in A compartments (red), across A and B compartments (purple) and in B compartments (blue). **e**, Overall inter-TAD distance of chr22 in cells with GFP and CHD7 overexpression. **f**, Radii of gyration of chr22 in cells with GFP and CHD7 overexpression. P values in **c**, **d** and **f** were calculated by two-sided Wilcoxon rank sum test. P value in **e** were calculated by two-sided Wilcoxon signed rank test.

**Extended Data Fig. 5.**
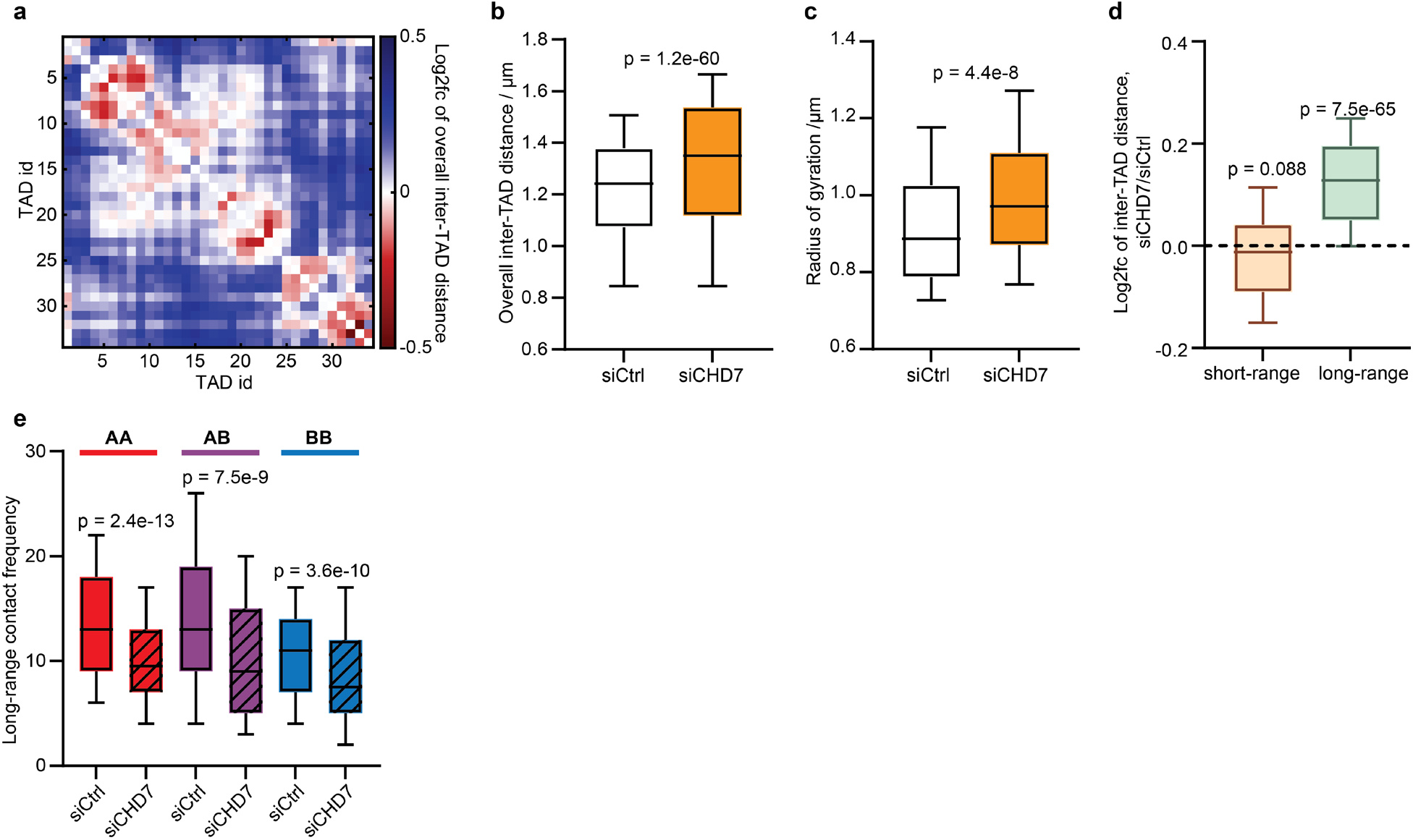
Validation of CHD7 perturbation phenotypes in a different cell background and genomic context. **a**, Log2 fold change matrix of overall inter-TAD distance of chr21 between siCHD7 and siCtrl in hTERT RPE-1 cells. **b**, Overall inter-TAD distance of chr21 in siCtrl and siCHD7 cells. **c**, Radii of gyration of chr21 in siCtrl and siCHD7 cells. **d**, Log2 fold change of short-range and long-range inter-TAD distances between siCHD7 and siCtrl. **e**, Compartmental contact frequencies in A compartments (red), across A and B compartments (purple) and in B compartments (blue) of chr21 in siCtrl and siCHD7 cells. P values in **b** and **d** were calculated by two-sided Wilcoxon signed rank test. P values in **c** and **e** were calculated by two-sided Wilcoxon rank sum test. Number of traces analyzed: 904 (siCtrl) and 210 (siCHD7).

**Extended Data Fig. 6.**
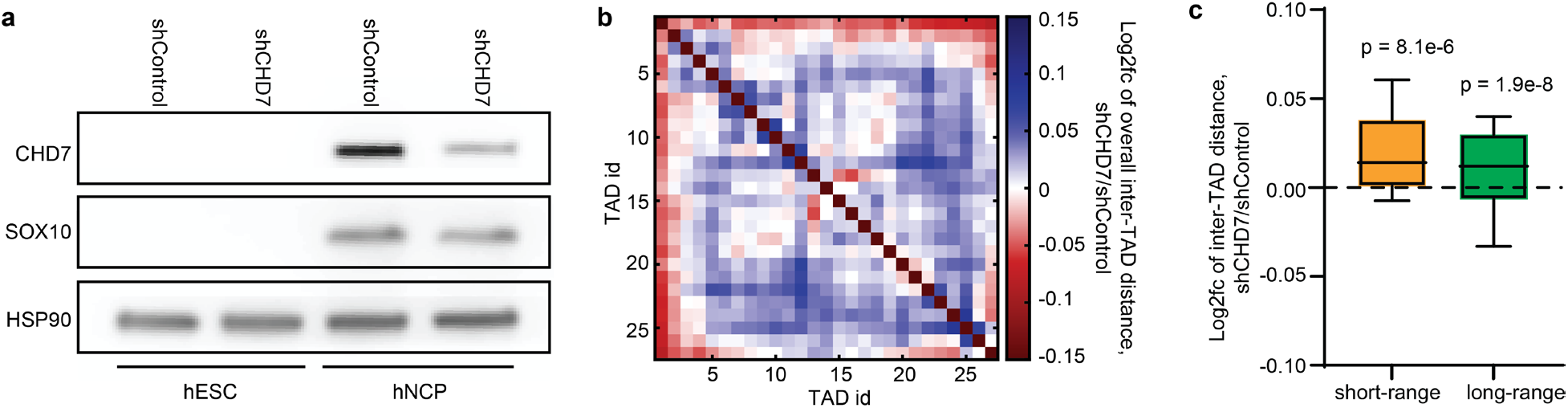
Validation of CHD7’s long range chromatin compaction function in neural crest cells. **a**, Western blot of shControl- and shCHD7-transduced human embryonic stem cells (hESC) and human neural crest progenitors (hNCP). Top: anti-CHD7 antibody; middle: anti-SOX10 antibody; bottom: anti-HSP90 antibody. CHD7 increased upon neural crest induction, and reduced in shCHD7 hNCP cells compared to shControl. Sox10, the neural crest marker, was expressed at similar levels in shControl and shCHD7 hNCP cells. HSP90 is a loading control. **b**, Log2 fold change of overall inter-TAD distance of chr22 between shCHD7 and shControl hNCP cells. Number of traces analyzed: 4,657 (shControl) and 2,796 (shCHD7). **c**, Log2 fold change of short-range and long-range inter-TAD distances of chr22 between shCHD7 and shContrl hNCP cells. P values were calculated by two-sided Wilcoxon signed rank test.

**Extended Data Fig. 7.**
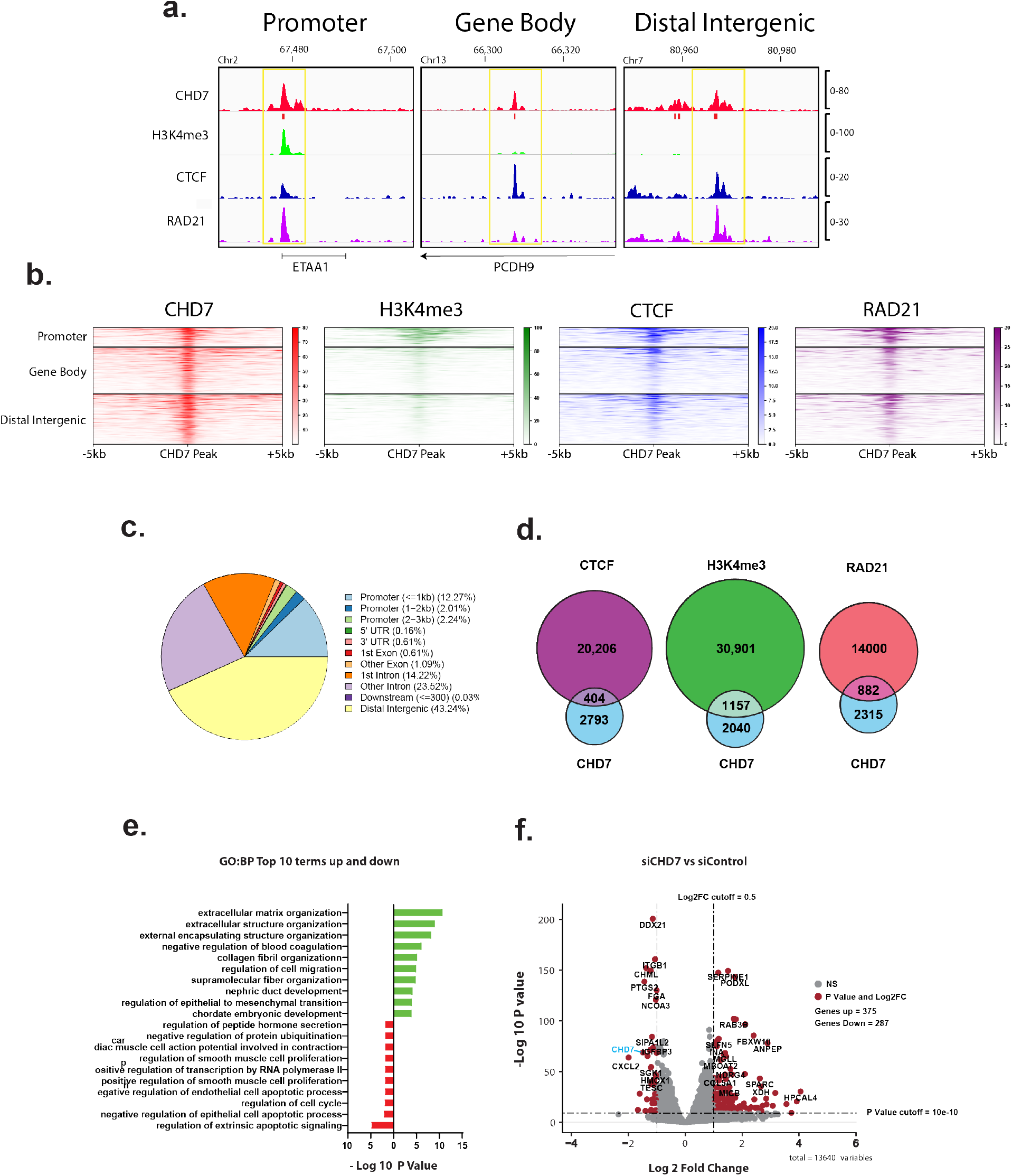
CHD7 binds diverse genomic regions. **a**, Example tracks of CUT&RUN peak profiles of CHD7 and other proteins/epigenetic mark over different genomic regions. **b**, Heat map of other proteins/epigenetic mark localized to CHD7 peaks by CUT&RUN. **c**, Peak annotation for all CHD7 CUT&RUN peaks. **d**, Overlap of CUT&RUN peaks of CTCF, RAD21, and H3K4me3 with CHD7 peaks. **e**, Top 10 gene ontology terms up and down in siCHD7 cells versus siControl cells based on bulk RNA-seq analyses. Gene ontology was performed using Enrichr. **f**, Volcano plot of RNA-seq comparing siCHD7 and siControl cells (siCHD7/siControl). Top differentially expressed genes are displayed on the graph as labels. CHD7 is highlighted and is a top differentially downregulated gene in the siCHD7 cells, validating the knockdown.

**Extended Data Fig. 8.**
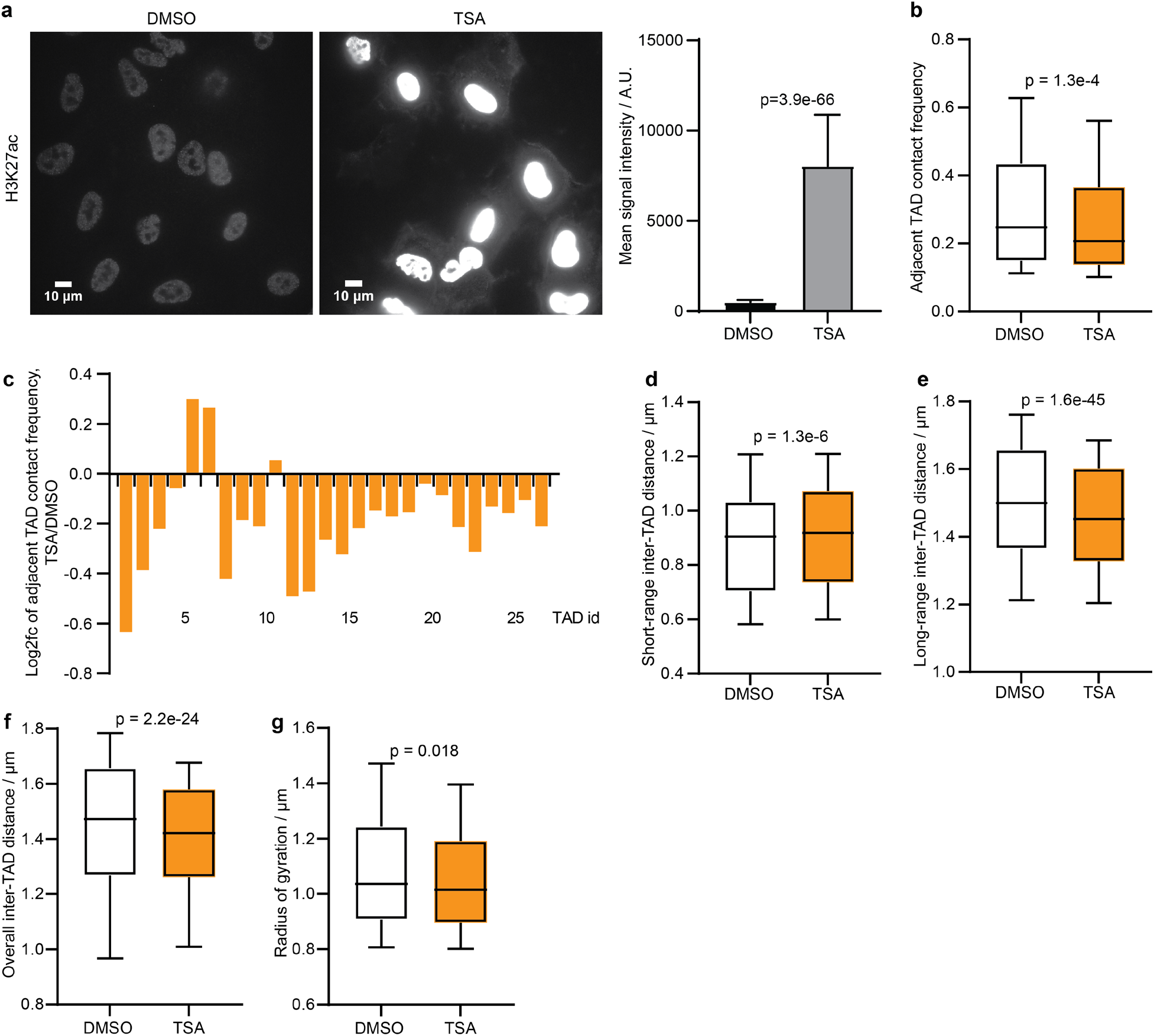
HDAC inhibition causes short-range chromatin decompaction and long-range chromatin compaction. **a**, H3K27ac IF measurements of DMSO-treated control (left) and TSA-treated (middle) A549 cells, and the quantification of mean signal intensity (right). The error bars represent standard deviations. Number of nuclei analyzed: 103 (DMSO) and 85 (TSA). **b**, Adjacent TAD contact frequency of DMSO- and TSA-treated cells. **c**, Log2 fold change of contact frequency between each pair of adjacent TADs along chr22 of TSA-treated cells compared to DMSO-treated cells. **d**, Short-range inter-TAD distance along chr22 of DMSO- and TSA-treated cells. **e**, Long-range inter-TAD distance along chr22 of DMSO- and TSA-treated cells. **f**, Overall inter-TAD distance of chr22 of DMSO- and TSA-treated cells. **g**, Radii of gyration of chr22 in DMSO- and TSA-treated cells. P value in **a** was calculated by two-tailed unpaired t test. P values in **b** and **d**-**f** were calculated by two-sided Wilcoxon signed rank test. P value in **g** was calculated by two-sided Wilcoxon rank sum test.

**Extended Data Fig. 9.**
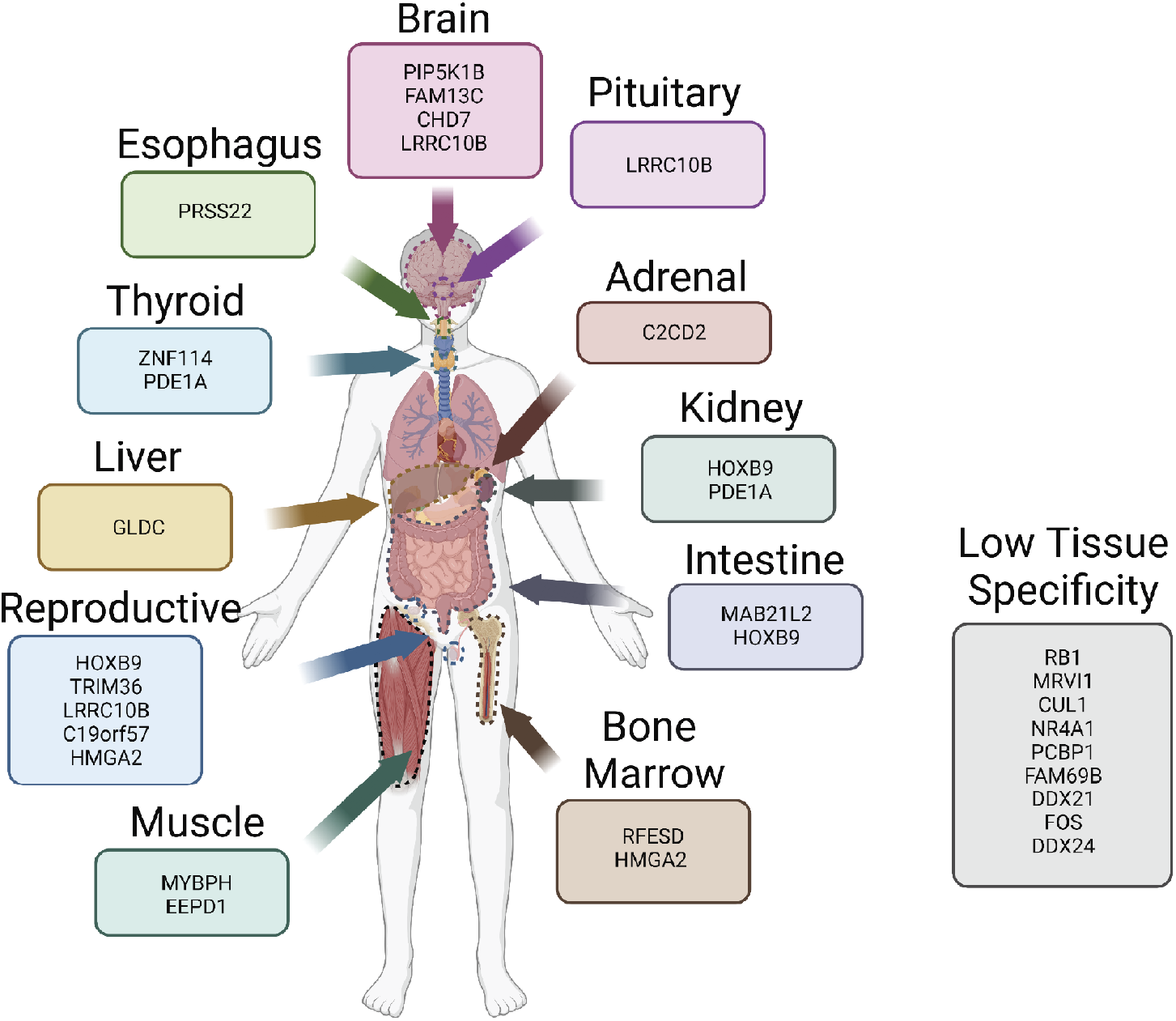
Tissue specific expression of top hits based on bulk RNA sequencing results from the Human Protein Atlas. This figure was created with Biorender.

## Data availability

All sequencing data (genomic DNA sequencing of the CRISPR screen cell library, RNA-seq, and CUT&RUN data) and analyzed imaging data generated for this study are available for download at https://campuspress.yale.edu/wanglab/BARCFISH. All sequencing data will also be available at the Gene Expression Omnibus (GEO) upon publication. Raw imaging data are available from the corresponding author upon request and are not deposited online due to prohibitively large size.

## Code availability

All original code generated for this study are available for download at https://campuspress.yale.edu/wanglab/BARCFISH, and will also be available on GitHub upon publication. Open-source codes for imaging data collection are available at https://github.com/ZhuangLab/storm-control.

## Acknowledgments

We thank Dr. Antonio Giraldez, Dr. Monkol Lek, and members of the Wang lab for helpful discussions. We thank members of the Yale Molecular Diagnostics Laboratory, Yale Center for Genome Analysis, the Keck DNA Sequencing Facility and the Yale Flow Cytometry Facility for their help. S. W. was partly supported by NIH (UG3CA268202, U01DA052775, R01HG011245, R33CA251037, DP2GM137414) and Pershing Square Sohn Cancer Research Alliance. B.Y., M.H. and Y.C. were in part supported by the China Scholarship Council. T.B.J. was in part supported by 5T32GM007205. J.S.D.R. was supported by NIH Predoctoral Training Grant (2T32GM007499). C.Z. was in part supported by NIH R35GM133712. This work was in part supported by NIH Director’s New Innovator Award (DP2GM137414) and Pershing Square Sohn Cancer Research Alliance.

## Author contributions

S.W. conceived the study. Y.C., T.Y., B.Y., M.H., T.B.J., R.Y., J.S.D.R., and S.W. performed experiments. Y.C., T.Y., M.H., B.Y., T.B.J., R.Y., Z.M., S.J., C.Z. and S.W. analyzed data. Y.C., T.Y., M.H., B.Y., T.B.J., Z.M. C.Z. and S.W. wrote the manuscript with inputs from all authors.

## Supplementary Information

### Methods

#### Barcode and probe sequence design

##### Design of BARC-FISH barcodes, linear and padlock probes

We first designed 60 barcode-targeting regions of linear and padlock probes using 46 published 20-nt sequences that only contain A, T and C^66, 87^ and 14 newly designed sequences. The 14 extra sequences were generated using a previously introduced orthogonal 25-nt oligonucleotide dataset^88^, by trimming off 5 consecutive nucleotides on the 5’ or 3’ end of the 25-nt sequences, and only sequences that met the following three criteria were retained: 1) the sequence only contains A, T and C; 2) the sequence does not contain 4 or more consecutive C’s; and 3) the percentage of C is between 40-45%. The newly designed 20-nt sequences were pooled with the 46 published sequences, and all these 20-nt sequences were BLASTed among themselves to ensure they were orthogonal to each other^89^. Finally, we BLASTed all possible pairs of concatenated 20-nt sequences separated by a G spacer against human genome and transcriptome to ensure they were orthogonal to endogenous nucleotide sequences^89^. We then grouped the 60 sequences into 30 pairs, and reverse-complemented the sequences to generate the DNA segments of barcode digits, each of which is 41-nt long (two 20-nt sequences with a 1-nt spacer). We adopted the backbone sequences of linear and padlock probes from a previous work^51^ to generate the full-length linear and padlock probe sequences, with the following modification: The backbone of padlock probes of all value 0 sequences for all barcode digits carried a 20-nt secondary probe targeting region. For value 1 and 2 sequences of the barcode digits, the 20-nt region of the padlock probe that binds to the barcode RNA serves as the secondary probe targeting region. 27 secondary probe sequences were previously reported^66, 90^. The other 3 secondary probe sequences were designed with a previously introduced procedure^91^. The linear and padlock probes, and secondary probes labeled with ATTO565, Alexa Fluor 647 and Alexa Fluor 750 dyes were individually ordered from Integrated DNA Technologies, Inc (IDT). A list of DNA barcode segment sequences and their corresponding linear, padlock and secondary probe sequences are included in Supplementary Table 3.

##### BARC-FISH helper probe design

To improve the hybridization efficiency of the linear and padlock probes, we designed helper probes targeting the flanking regions of the barcode RNA, which could help open potential secondary structures and linearize the RNA molecules. To design the helper probes, we adapted the algorithm of a published probe designing tool ProbeDealer^92^. Briefly, three RNA regions were used as the input targeting regions for ProbeDealer: an 888-nt region upstream of the protospacer, a 108-nt region between the protospacer and barcode, and a 310-nt region downstream of a unique molecular identifier region (for the region layout, refer to later sections on plasmid library construction). The probes were generated with the following constraints: The probe length was 30-nt, with at least 1-nt spacing between adjacent probes; the allowed melting temperature range was 66-100℃; the allowed GC content range was 30-90%; the melting temperature of internal secondary structure of each probe was no greater than 76℃; the melting temperature of cross-hybridization regions among the probes was no greater than 72℃; and the probes containing consecutive repeat of more than five identical nucleotides were excluded. The probe sequences were further BLASTed against human genome and transcriptome to ensure the hybridization specificity^89^. Probes with alignments to either human genome or transcriptome were excluded from the selection. In total, 36 helper probes were generated and ordered though IDT. The sequences of the helper probes are listed in Supplementary Table 4.

##### Chromatin tracing primary probes

We used the primary probes of human chr21 and chr22 from a previous publication^50^ with slight modifications on secondary probe binding regions. Specifically, the probes consisted of the following segments from the 5’ to 3’: 1) a 20-nt priming regions at 5’ end, 2) a secondary probe binding region of 17-19 nt, 3) a 30-nt primary binding region complementary to genomic DNA, 4) a same secondary probe binding region of 17-19 nt, and 5) a 20-nt priming region at 3’ end. The secondary probe binding regions were generated by trimming the 5’ or 3’ ends of previously published 30-nt secondary probe binding regions^50^, so that the Tm of the trimmed secondary probe binding regions became 50-57°C.

#### Barcode plasmid library construction

The DNA sequences of the 10-digit barcodes each comprised ten 41-nt digit segments, with a nucleotide ‘C’ separating adjacent digits. Each digit contained one of three different 41-nt sequences, representing the three different values (0, 1, 2) of the ternary digit. The values from the 10 digits were randomly assembled. Therefore, theoretically 3^10^ or 59,049 possible unique combinatorial barcodes could be generated. To assemble the barcode library, we used a pooled barcode cloning strategy (Extended Data Fig. 2a) modified from approaches previously described^49, 78^: First, the 10 digits were divided into two fragments. Overlapping oligos encoding and bridging digit 1-5 were mixed to a final concentration of 100 nM each, and the mixture was subjected to phosphorylation and ligation. Oligos encoding digit 6-10 were treated in the same way in parallel. Then, the two 5-digit fragments were gel-purified and assembled into full-length barcodes by overlapping PCR. Next, the full-length barcodes were amplified by limited-cycle PCR. In this step, each barcode molecule was paired with a unique molecular identifier (UMI), a 20-bp random sequence, at the 3’ end for later next-generation sequencing (NGS), and overhangs at both ends for Gibson Assembly. The PCR products were gel-purified and then assembled into a plasmid backbone digested with MluI (New England BioLabs, R3198S) and SpeI (New England BioLabs, R3133S), through Gibson Assembly (New England Biolabs, E2621L). The backbone plasmid was modified from plasmid LentiGuide-Puro (Addgene, 52963), with an addition of a CMV promoter, an EGFP gene and two restriction sites, MluI and SpeI. Lastly, the Gibson reaction products were purified by isopropanol precipitation and electroporated into Endura electrocompetent cells (Lucigen, 60242-2). The transformed bacteria were spread onto LB agar plates containing 100 µg/mL ampicillin and incubated overnight at 37 ℃. On the next day, ∼70 million colony forming units grown on the plates were collected and subjected to maxi-prep (Qiagen, 12362) to extract the barcode plasmid library. The barcode plasmid library was then used to generate CRISPR screen plasmid library. All oligos and primers were ordered from IDT. The oligo and primer sequences for constructing the barcode plasmid library are provided in Supplementary Table 5. The modified backbone sequence with annotations is included in Supplementary File S1. An example barcode plasmid sequence with annotations is included in Supplementary File S2.

#### sgRNA plasmid library construction

To construct the sgRNA plasmid library, we selected protospacer sequences from a previous publication^93^. The sgRNA fragments were amplified from a CustomArray oligo pool via limited-cycle PCR. The backbone was digested with FastDigest Esp3I (Thermo Fisher Scientific, FD0454) together with Alkaline Phosphatase (Thermo Fisher Scientific, EF0654) overnight at 37 ℃. The backbone plasmid was modified from plasmid LentiGuide-Puro with an addition of a CMV promoter and an EGFP gene. The sgRNA fragments and digested backbone were gel-purified and assembled through Gibson Assembly. The Gibson reaction products were purified by isopropanol precipitation and the electroporated into Endura electrocompetent cells. The cells were plated onto LB agar plates containing 100 µg/mL ampicillin and incubated at 37℃ overnight. About 2.5 million colony forming units were collected and inoculated into 200 mL LB liquid medium containing 100 µg/mL ampicillin, and grown overnight to amplify the sgRNA plasmid library. The sgRNA plasmid library was maxi-prepped from the overnight culture and used for the construction of CRISPR screen plasmid library. All primers were ordered from IDT. The protospacer sequences, ordered CustomArray oligo sequences and primer sequences are provided in Supplementary Table 6. The modified backbone sequence with annotations is included in Supplementary File S3. An example sgRNA plasmid sequence with annotations is included in Supplementary File S4.

#### CRISPR screen plasmid library construction

We adopted the cloning strategies from previous publications^49, 94^ to construct the CRISPR screen plasmid library (Extended Data Fig. 2a). In brief, sgRNA, barcode and backbone were assembled through Gibson Assembly. The barcode fragments were amplified by limited-cycle PCR from the pre-made barcode plasmid library. Similarly, the sgRNA fragments were PCR-amplified from the pre-made sgRNA plasmid library. The backbone plasmid was modified from plasmid CROPseq-Guide-Puro (Addgene, 86708), with an addition of a CMV promotor and removal of the gRNA scaffold. The modified plasmid was restriction digested with FastDigest Esp3I overnight at 37℃. The restriction reaction product was treated with Alkaline Phosphatase to remove the phosphate groups from the linearized backbone. The DNA fragments of sgRNA, barcode and backbone were gel purified and mixed for Gibson Assembly. The Gibson reaction products were purified by isopropanol precipitation and electroporated into Endura electrocompetent cells. Electroporated cells were grown overnight at 37℃ on LB agar plates containing 100 µg/mL ampicillin. To restrict the number of barcodes paired with each sgRNA and allow unique projection from each barcode to a single sgRNA, we used a bottlenecking strategy: Only 4,000∼5,000 bacterial colonies were collected from the plates and cultured in 200 mL LB liquid medium overnight at 37℃ with shaking at 225 rpm. The CRISPR screen plasmid library was then extracted and purified by maxi-prep. All primer sequences are provided in Supplementary Table 7. The modified backbone sequence with annotations is included in Supplementary File S5. An example CRISPR screen plasmid sequence with annotations is included in Supplementary File S6.

#### Plasmid construction for CHD7 and GFP overexpression

The GFP lentiviral overexpression vector, pLenti-GFP, was purchased from OriGene (OriGene, PS100093). To construct the CHD7 lentiviral overexpression construct, we cloned the human CHD7 open reading frame (ORF) into the same vector upstream of GFP, together with a P2A sequence placed between CHD7 and GFP for self-cleavage. Specifically, the CHD7 ORF was PCR amplified from a plasmid (Addgene, 89460) using Phusion High-Fidelity Master Mix (NEB, M0531L) following the manufacturer’s instructions. The P2A sequence was added to the 3’ end via reverse primer. Then, DpnI (NEB, R0176S) was directly added to the PCR mixture and incubated at 37℃ for 1 hour to digest the plasmid template. The PCR products of correct size were confirmed by agarose gel electrophoresis and gel purified. To prepare the backbone, pLenti-GFP was digested by restriction enzymes AsiSI (NEB, R0630S) and MluI-HF (NEB, R3198S), and dephosphorylated by Shrimp Alkaline Phosphatase (NEB, M0371S) at 37℃ for 3.5 hours. The resulting backbone was gel purified, mixed with the purified PCR products, and subjected to Gibson Assembly using NEBuilder HiFi DNA Assembly Master Mix (NEB, E2621L) following the manufacturer’s instructions. The reaction mixture was then column purified and transformed into chemically competent E. coli Stbl3 (ThermoFisher, C737303) following the manufacturer’s instructions. The transformed cells were plated onto LB agar plates containing 34 µg/mL chloramphenicol and incubated overnight at 37℃. Several clones were picked and individually inoculated into LB liquid medium overnight at 37℃, ∼225 rpm. The plasmids of individual clones were extracted using QIAprep Spin Miniprep Kit (Qiagen, 27106) following the manufacturer’s instructions, and the correct sequence was confirmed by Sanger sequencing. The primer sequences for cloning and Sanger sequencing are provided in Supplementary Table 8. The plasmid sequence with annotations of the reconstructed CHD7 lentiviral overexpression vector, pLenti-CHD7, is included in Supplementary File S7. For the construction of both CHD7-ΔBRK overexpression construct and CHD7-K999R overexpression construct, pLenti-CHD7 was modified by a three-fragment Gibson Assembly. For CHD7-ΔBRK overexpression construct, the first fragment was generated by digestion of pLenti-CHD7 by restriction enzymes ScaI-HF (NEB, R3122S) and NheI-HF (NEB, R3131S) followed by gel purification. For CHD7-K999R overexpression construct, the first fragment was generated by digestion of pLenti-CHD7 by restriction enzymes SphI-HF (NEB, R3182S) followed by gel purification. Other two fragments were PCR amplified from pLenti-CHD7 using Phusion High-Fidelity Master Mix (NEB, M0531L) following the manufacturer’s instructions. The PCR products of correct size were confirmed by agarose gel electrophoresis and gel purified. Next, Gibson Assembly, transformation and plasmids extraction were performed as mentioned above. The primer sequences for cloning and Sanger sequencing are provided in Supplementary Table 8. The plasmid sequence with annotations of pLenti-CHD7-ΔBRK is included in Supplementary File S8. The plasmid sequence with annotations of pLenti-CHD7-K999R is included in Supplementary File S9.

#### Lentivirus production and cell line construction

##### Lentivirus production

The HEK-293FT cells (Thermo Fisher Scientific, R70007) were cultured to be 70-90% confluent at the point of transfection to produce lentiviruses. The Cas9 plasmid lentiCas9-Blast (Addgene, 52962), the GFP overexpression plasmid pLenti-GFP, the CHD7 overexpression plasmid pLenti-CHD7, or the CRISPR screen plasmid library was mixed with helper plasmids psPAX2 (Addgene, 12260) and pVSV-G (Addgene, 138479) together with Lipofectamine 2000 (Thermo Fisher Scientific, 11668019) according to the manufacturer’s instructions. The mixture was then added into the cell culture and incubated for two days. After the lentiviral transfection, the lentivirus supernatant was collected from the cell culture and filtered through a 0.45 µm strainer (Millipore, SLHAR33SS) to remove cell debris. The filtered lentivirus supernatant was concentrated using Amicon Ultra-15 Centrifugal Filter Unit (Millipore, UFC910024) following the manufacturer’s instructions. The concentrated lentivirus supernatant was aliquoted and stored at -80℃.

##### Generation of clonal A549-Cas9 cells

A549 cells (ATCC, CCL-185) were infected with lentivirus generated from the lentiCas9-Blast plasmid. Cells were under Blasticidin selection at 10 µg/mL until all untransduced cells were killed. The Blasticidin-selected, polyclonal cells were sorted into single cells by flow cytometry and plated on 96-well plates. The single cells were clonally expanded and analyzed by immunofluorescence to verify Cas9 expression. The clonal selection was important to reduce the heterogeneity in the cell background in the screen.

##### Generation of BARC-FISH CRISPR screen cell library

Clonal A549-Cas9 cells were cultured to 60-80% confluency and transduced with the CRISPR screen lentivirus supernatant at an MOI < 0.3 to ensure that most cells received only one genetic perturbation. Two days after the lentiviral infection, Puromycin (Thermo Fisher Scientific, A1113803) was added into the media at 3 µg/mL to select the cells with resistance. Cells were under Puromycin selection for 10-12 days until the cell library was established. The cell library was then aliquoted into frozen stocks and stored in liquid nitrogen.

##### Generation of CHD7 and GFP OE cell line

Clonal A549-Cas9 cells were infected with the concentrated lentivirus produced from the overexpression plasmid pLenti-GFP or pLenti-CHD7. Puromycin selection started two days after lentiviral infection at a concentration of 3 µg/mL in growth media, and lasted for 5-6 days until the stable cell lines were established.

##### Human ESC culture

H1 human ESC lines were culture in mTeSR1 (Stem Cell Technologies) in feeder free conditions on 1x hESC certified matrigel (Corning 354277) with daily media changes for the duration of culture. Cells were passaged weekly with dispase maintaining ESCs in colonies (Stem Cell Technologies 07913). Cells were weeded manually whenever differentiation was observed.

When ready to differentiate cells, ESCs were detached and made into a single cell suspension using Accutase (Stem cell technologies 07920).

##### Generation of CHD7 knockdown H1 hESC line

With the Yale Functional Genomics Core, lentiviruses were produced by co-transfecting HEK293T cells with packaging vectors psPAX2 (Addgene plasmid #12260) and pMD2.G (Addgene plasmid #12259) together with lentiviral transfer constructs. Viral supernatant was collected 48h and 72h after transfection and filtered with a 0.45-µm filter. Viral supernatant, either Sigma Mission SHC002 nontargeting shRNA, to be referred to as shControl, or shCHD7 (Sigma Mission shRNA TRCN0000016408) was diluted 1:5 into mTeSR1 and plated onto cells for 24 hours. After infection, the viral media was replaced with mTeSR1 for an additional 24 hours before beginning selection. Cells were selected in 10 µg/mL puromycin with daily media changes for 7 days, at which point the cells were taken off selection. Prior kill curves in house have determined the kill time of 10 µg/mL puromycin in wild type cells to be 2 days. A well of uninfected cells was included on each plate as a control to verify 100% cell death without puromycin resistance. Cells were passaged or frozen after expansion following selection. Cells were frozen in 40% Knockout serum replacement (Thermofisher 10828028) 10% DMSO and 50% mTeSR1 and were slow frozen overnight in an isopropanol bath.

##### Generation of Neural Crest

Knockdown and control transfected hESCs were differentiated to neural crest using the STEMdiff Neural Crest Differentiation Kit (Stem cell technologies 08610) exactly according to manufacturer’s specifications. In brief, hESCs were dissociated into single cells with Accustase and seeded as a single cell suspension in complete neural crest media supplemented with 10 µM Y-27632 (Dihydrochloride) (Stem cell technologies 72304). Neural crest media was changed daily for 6 days. Afterwards, cells were passaged with accutase onto coverslips coated with Matrigel. These coverslips were then fixed and treated as described for chromatin tracing and IF 1-2 days after passaging onto coverslips and were not maintained long term but were generated fresh as needed from frozen hESC stocks.

#### Cell culture for cloning and imaging

A549 cells and the clonal A549-Cas9 cells (introduced in the section above, derived from the A549 cells) were maintained in F-12K media (Corning, 10-025-CV) supplemented with 10% FBS (VWR, 97068-091) and 1% Pen-Strep (Thermo Fisher Scientific, 15140122) at 37℃ in 5% CO_2_. hTERT RPE-1 cells (ATCC, CRL-4000) were maintained in DMEM:F-12 media (ATCC, 30-2006) supplemented with 10% FBS (VWR, 97068-091) and 1% Pen-Strep (Thermo Fisher Scientific, 15140122) at 37℃ in 5% CO_2._ To prepare imaging samples, coverslips (Bioptechs, 40-1313-03193) were disinfected by 15-min UV treatment on each side prior to use. To culture cells for conducting the screen, A549-Cas9 cells transduced with BARC-FISH screen library at the same passage were seeded onto the disinfected coverslips at 10% density in F-12K media supplemented with 15% FBS and 1% Pen-Strep and cultured for 6 days. The higher percentage of FBS in the media allowed the cells to produce more barcode RNA copies. The media was also supplemented with 3 µg/mL Puromycin and 10 µg/mL Blasticidin (Thermo Fisher Scientific, R21001) to ensure sgRNA and Cas9 expression. Media were refreshed every 3-4 days. For lentivirus production, HEK-293FT cells were cultured in DMEM medium supplemented with GlutaMAX-I (Thermo Fisher Scientific, 10569-010), 1× MEM Non-Essential Amino Acids (Thermo Fisher Scientific, 11140-050), 10% FBS and 1% Pen-Strep at 37℃ in 5% CO_2_, and were used within 10 passages. All cells were routinely tested of mycoplasma contamination via Yale Molecular Diagnostics Laboratory.

#### Detection of sgRNA-barcode associations in the cell library

##### Next-generation sequencing (NGS) library preparation for mapping sgRNA-barcode associations

To determine the sgRNA-barcode correspondence in the screen library, we PCR-amplified the cell genomic DNA for NGS. Because the total length of the sgRNA-barcode-UMI cassette is longer than the maximum sequencing length of NGS (300 bp paired-end sequencing on Illumina MiSeq), we generated two sequencing libraries: an sgRNA-barcode-UMI library and a barcode-UMI library. The sgRNA-UMI correspondence can be determined by the sgRNA-barcode-UMI library and the barcode-UMI correspondence can be determined by the barcode-UMI library. Then the sgRNA-barcode correspondence can be determined through their common UMI associated with both the sgRNA and the barcode on the same molecule. Briefly, the genomic DNA was extracted from the CRISPR screen cell library using Zymo Quick-DNA Miniprep Kit (Zymo Research, D4068) following the manufacturer’s protocol. To achieve a sequencing coverage of >500, more than 2.5 million cells were harvested to cover the ∼5,000 barcode varieties. We amplified all harvested genomic DNA and limited the number of amplification cycles to 20-23 to reduce amplification bias. For PCR amplification, each 50 µL reaction mixture was composed of 1 µg of genomic DNA, 500 nM of each forward and reverse primers, 5% DMSO and 1× NEBNext High-Fidelity PCR master mix (New England BioLabs, M0541L) following the manufacturer’s instructions. The PCR primers contained flow-cell binding sequence, sequencing index, sequencing primer binding region, and a 9-13 bp random sequence that improved the library diversity. After the reactions were completed, the PCR products were purified and concentrated using Zymo DNA Clean & Concentrator (Zymo Research, D4030) following the manufacturer’s instructions. Lastly, the concentrated products were run on a 2% agarose gel to validate the correct DNA amplicon size. The DNA library amplicons with the correct size were then gel-purified and eluted with elution buffer (10 mM Tris-HCl, pH 8.5) or MilliQ water. The DNA libraries were sequenced on Illumina MiSeq system in 2×300 bp format. The PCR primers were all purchased from IDT and the sequences are provided in Supplementary Table 9.

##### sgRNA-barcode NGS analysis

As described above, we generated two sequencing libraries, a sgRNA-barcode-UMI library and a barcode-UMI library, and determined the sgRNA-barcode correspondence by the common UMI sequence. Briefly, the sequencing reads were filtered by read length and quality score. From the sgRNA-barcode-UMI library, protospacer and UMI sequences were extracted from the reads, and the protospacer-UMI look-up table was generated accordingly. The protospacer sequences were aligned with the sequences in the pre-designed sgRNA oligo library such that the reads with improper protospacers were removed. From the barcode-UMI library, the UMI sequences were extracted and compared to the UMI found in the sgRNA-barcode-UMI library. Only reads with common UMIs were retained for further analysis. Because the read length from a single end did not cover the full length of the barcode, the sequencing read from each end only contained a partial barcode sequence. The digit values from the partial barcode sequences were extracted by BLAST^89^ against the pre-designed digit sequences. The two decoded partial codes assigned to the two ends of the same sequence were then assembled into one full-length complete code by the overlapping region of the two partial codes. The partial codes that failed to overlap were excluded. The UMI-barcode lookup table was then established. The two look-up tables were merged to determine the sgRNA-barcode correspondence (codebook) through the shared UMIs. This codebook allowed analyses of the percentage of good codes (codes uniquely associated with one sgRNA) versus bad codes (codes associated with multiple sgRNAs) (Extended Data Fig. 1d). The final codebook containing both the good codes that each uniquely associate with one sgRNA and the bad codes that each have multiple sgRNA projections is listed in Supplementary Table 10.

#### Validation of CRISPR knockout efficiency

To validate the CRISPR knockout efficiency, we constructed individual CRISPR knockout cell lines for selected sgRNAs and utilized NGS of genomic DNA to analyze the frameshift indel mutations. The cloning procedure of individual sgRNA plasmids was similar to previously described in “sgRNA plasmid library construction”, except that we directly ordered double-stranded sgRNA fragments from IDT for Gibson Assembly. Cell line generation procedure was identical to “Generation of BARC-FISH CRISPR screen cell library”. We then performed genomic DNA extraction and PCR amplification following a procedure similar to previously described in “Next-generation sequencing (NGS) library preparation for mapping sgRNA-barcode associations”. The amplified region was 250-450 bp centered around the sgRNA targeting site. The PCR amplicons were sequenced on Illumina MiSeq system in 2×250 bp format, achieving ∼50,000 reads per sgRNA. The indel rate analysis was performed and delivered by Genewiz.

#### Probe synthesis

The template oligo pool for synthesizing chromatin tracing primary probes was ordered from CustomArray, GenScript. We adopted a previously described probe synthesis workflow^90, 91^ of limited-cycle PCR, *in vitro* transcription, reverse transcription, alkaline hydrolysis and column purification. The PCR primers and reverse transcription primers were ordered from IDT. The sequences of the template oligo library and primers are included in Supplementary Table 11.

#### Imaging sample preparation

##### Geminin antibody staining

All steps were performed at room temperature unless otherwise described. Cells grown on coverslips were briefly rinsed with DPBS after media removal, and then fixed in 4% paraformaldehyde (PFA) (Electron Microscopy Sciences, 15710) diluted in DPBS for 10 minutes followed by three DPBS washes. Cells were then permeabilized with 0.5% v/v Triton X-100 in DPBS for 10 minutes followed by three DPBS washes. After permeabilization, cells were blocked in blocking buffer (1% w/v bovine serum albumin (Sigma-Aldrich, A9647), 0.1% v/v Tween-20 in DPBS) supplemented with 0.1% v/v murine RNase inhibitor (MRI) (New England Biolabs, M0314L) for 30 minutes, followed by primary antibody incubation for 1 hour with 1:100 diluted anti-Geminin antibody (Abcam, ab195047) in blocking buffer supplemented with 1% v/v MRI. Unbound primary antibody was washed three times with 0.1% v/v Tween-20 in DPBS (DPBSTw), 5 minutes each, followed by a 1-hour incubation with 1:1000 diluted Alexa Fluor 488-labeled secondary antibody (Invitrogen, A11034) in blocking buffer supplemented with 1% v/v MRI. Starting from secondary antibody incubation, samples were protected from light during all steps. Excessive secondary antibody was washed off by washing three times, 5 minutes each with DPBSTw, and samples were post-fixed in 4% PFA in DPBS for 10 minutes followed by 3 DPBS washes. Post-fixed samples were next used for the BARC-FISH procedure in the screen experiments and for chromatin tracing primary probe hybridization in the validation experiments.

##### BARC-FISH

Helper probes and BARC-FISH linear and 5’-phosphorylated padlock probes were ordered from IDT. A total of 36 helper probes were pooled in an equimolar manner to a concentration of 5.6 µM for each probe. A total of 60 probes for BARC-FISH (30 linear probes and 30 padlock probes) were pooled in an equimolar manner to a concentration of 1.67 µM for each probe. Immediately prior to use, the pooled BARC-FISH probes were heat-denatured at 90°C for 2-5 minutes, and then cooled down to room temperature. Samples with Geminin staining were pre-hybridized in 2× saline-sodium citrate buffer (2×SSC) containing 20% v/v formamide and 0.1% v/v Tween-20 for 10 minutes, and then 50 μL of primary hybridization buffer was added to each sample, which contained 20% v/v formamide, 0.1 mg/mL salmon sperm DNA (Invitrogen, 15632-011), 800 nM each of the 60 BARC-FISH probes, 100 nM each of the 36 helper probes and 1% v/v MRI in 2×SSC. Samples were then incubated at 37°C for 16-20 hours. To remove excessive primary probes, samples were washed twice in 2×SSC containing 40% v/v formamide and 0.1% v/v Tween-20, 15 minutes each, followed by a third wash for 20 minutes at 37°C in buffer containing 4×SSC, 1×DPBS, 0.1% v/v Tween-20 and 0.1% v/v MRI. Samples were then briefly rinsed twice with DPBSTw prior to the T4 ligation step. The T4 ligation mixture contained 0.2 mg/mL BSA (New England Biolabs, B9000S), 1 mM extra-supplemented dithiothreitol (DTT) (Thermo Scientific, R0861), 1 mM extra-supplemented ATP (Thermo Scientific, R0441), 0.5 U/μL T4 DNA ligase (Thermo Scientific, EL0014) and 1% v/v MRI in 1× T4 ligase buffer (Thermo Scientific, EL0014). To perform the T4 ligation step, samples were incubated with 50 μL of T4 ligation mixture for 2 hours, followed by two washes in DPBSTw. Samples were then incubated in 30°C water bath for 5 hours with 50 μL of RCA mixture that contained 250 μM dNTP (New England Biolabs, N0447L), 1 mM extra-supplemented DTT, 0.2 mg/mL BSA, 1 U/μL phi29 enzyme (Thermo Scientific, EP0092) and 1% v/v MRI in 1× phi29 buffer (Thermo Scientific, EP0092). After the RCA step, samples were washed in DPBSTw, post-fixed with 4% PFA in DPBS for 30 minutes, and washed with DPBS. Post-BARC-FISH samples were used for the chromatin tracing primary probe hybridization in the screen experiments.

##### Chromatin tracing primary probe hybridization

Post-BARC-FISH samples (for screen experiments) or post-Geminin staining samples (for validation experiments) were briefly rinsed in DPBS, and then incubated with 0.1M HCl diluted in water for 5 minutes. After two DPBS washes, samples were treated with 0.1 mg/mL RNase-A (AB12023-00100) diluted in DPBS for 45 minutes at 37°C, followed by two DPBS washes and one 2×SSC wash. Subsequently, samples were pre-hybridized in 2×SSC containing 50% v/v formamide and 0.1% v/v Tween-20 for 30 minutes and carefully dried by dipping on tissue paper to remove excessive pre-hybridization buffer. 25 <u>μ</u>L of hybridization buffer containing 50% v/v formamide, 20% v/v dextran sulfate (Millipore, S4030) and 4 μM (total concentration) chromatin tracing primary probes in 2×SSC was applied to a glass slide, and the sample coverslip was carefully flipped and placed on top so that the glass slide and coverslip “sandwiched” the hybridization buffer, with the cells submerged into the hybridization buffer. The samples were then heat-denatured on an 86°C heat block (with a surface temperature of ∼80°C) for 3 minutes with the glass slide touching the heat block, and incubated in a humid chamber for 16-20 hours at 37°C. To remove excessive primary probes, samples were washed twice in a 60°C water bath with 0.1% v/v Tween-20 in 2×SSC, 15 minutes each, followed by a third 15-minute wash with 0.1% v/v Tween-20 in 2×SSC at room temperature. Samples were then briefly rinsed with 2×SSC, and 1:250,000 diluted yellow-green fiducial beads (Invitrogen, F8803) in 2×SSC were applied to the samples and incubated for 10 minutes. Excessive beads were removed by two 2×SSC washes. Post-hybridization samples were used for automated sequential imaging.

#### Automated sequential imaging

##### Imaging system setup

Four home-built fluorescence microscopes were used for image acquisition^90^. Each microscope had a Nikon Ti2-U body, a Nikon CFI Plan Apo Lambda 60× oil objective lens (NA 1.40) and an automated focus-lock system^90, 95^. One setup has identical lasers, light paths and filters as introduced previously^90^, and the image size was 1536×1536 pixels with a pixel size of 108 nm. For the other three setups, a Lumencor CELESTA light engine was used for illumination, with the following laser wavelengths: 405 nm, 477 nm, 546 nm, 638 nm and 749 nm. The 405 nm laser was used to excite and image the nuclear stain with diamidino-2-phenylindole (DAPI). The 477 nm laser was used to excite and image the yellow-green fiducial beads and the Geminin stain. The 546 nm laser was used to excite and image ATTO565 dye on secondary probes for BARC-FISH and chromatin tracing, and CF568 dye for total protein stain. The 638 nm laser was used to excite and image Alexa Fluor 647 dye on secondary probes for BARC-FISH and chromatin tracing. The 749 nm laser was used to excite and image Alexa Fluor 750 dye on secondary probes for BARC-FISH. A pentaband dichroic mirror supplied by Lumencor for the light engine was installed on the excitation path to direct the lasers to the sample, together with a ND0.6 neutral density filter to reduce the laser intensity. A pentaband emission filter supplied by Lumencor for the light engine and a Hamamatsu Orca Flash 4.0 V3 camera were each installed on the emission path. The image size was 2048×2048 pixels with a pixel size of 108 nm. A motorized x-y stage (SCAN IM 112×74, Marzhauser) was used to automatically image different fields of view (FOVs). Samples cultured on 40 mm #1.5 coverslips (Bioptechs, 40-1313-03193) were assembled in a Bioptech’s FCS2 flow chamber, mounted onto the microscope stage and connected with a previously described automated fluidic system^88, 89^ for liquid handling during sequential hybridization and imaging.

##### Chromatin tracing sequential imaging

We conducted 17 and 14 rounds of three-color imaging for chromatin tracing of chr21 and chr22, respectively. During each round of hybridization, the sample was first incubated with 20% v/v ethylene carbonate (EC) (Sigma-Aldrich, E26258) in 2×SSC containing two secondary probes (one labeled by ATTO565 and one labeled by Alexa Fluor 647) for 25 minutes. The final concentration of each secondary probe was 6 nM except the following exceptions for chr22 tracing to tune signal intensity differences: TAD4 (0.5 nM), TAD14 (12 nM), TAD18 (12 nM) and TAD26 (12 nM). In 6 initial screen replicates out of the 17 total replicates, we used a TAD4 probe concentration of 6 nM, which led to a bleedthrough of TAD4 signal (labeled with Alexa Fluor 647 dye) into the TAD18 fluorescent channel (546 nm laser channel). This bleedthrough issue was computationally addressed as described later. The dye-labeled secondary probe sequences were selected from a previous study^50^. The secondary probes were ordered from IDT and their sequences are listed in Supplementary Table 12. 2 mL of wash buffer containing 20% v/v EC in 2×SSC was flowed through the chamber to remove unbound secondary probes, followed by 2 mL anti-photobleaching oxygen-scavenging imaging buffer that contained 50 mM Tris-HCl pH 8.0, 5% w/v glucose, 2 mM Trolox (Sigma-Aldrich, 238813), 0.5 mg/mL glucose oxidase (Sigma-Aldrich, G2133) and 40 μg/mL catalase (Sigma-Aldrich, C30) in 2×SSC. During the overnight imaging process, the imaging buffer in the input tube was covered by a layer of mineral oil (Sigma-Aldrich, 330779-1L) to prevent oxidization. At each FOV, three z image stacks were taken sequentially with 638-nm, 546-nm and 477-nm laser illumination, with a step size of 200 nm, 0.4 s exposure time at each step, and a total z range of 7 μm. After all the FOVs were imaged for the current imaging round, the secondary probes were stripped off by slowly flowing 4 mL of 65% v/v formamide in 2×SSC for 10 minutes and incubating for additional 5 minutes. Extra formamide was then removed by flowing 2 mL of wash buffer, and the next round of hybridization and imaging followed. To correct for color shift between 546-nm and 638-nm lasers, a #1.5 coverslip was coated with 100-nm Tetraspeck beads (Invitrogen, T7279) diluted at 1:200 in the imaging buffer, and z-stack calibration images were taken using the two lasers^50, 90^.

##### BARC-FISH sequential imaging

For screen samples, we conducted 10 rounds of four-color imaging to detect the barcode after chromatin tracing. During each round, the sample was first incubated for 25 minutes with 20% v/v EC in 2×SSC containing three secondary probes that correspond to the three values in each barcode digit, and the three probes were labeled with ATTO565, Alexa Fluor 647 and Alexa Fluor 750, respectively. The concentrations of the Alexa Fluor 750- and Alexa Fluor 647-labeled probes were 3 nM each, and the concentrations of ATTO565-labeled probes were 6 nM each. Excessive probes were washed away by flowing 2 mL of wash buffer, followed by application of 2 mL imaging buffer. At each FOV, four z image stacks were taken sequentially with 749-nm, 638-nm, 546-nm and 477-nm lasers. Each z-stack had a step size of 1.5 μm, exposure time at each step of 0.4 s, and a total range of 9 μm. After each imaging round, 11 mL of 90% v/v formamide in DPBS was flowed through the sample chamber for 28 minutes and then let stand for 100s to remove the bound fluorescent probes. Excessive formamide was removed with 2 mL wash buffer.

##### DAPI and total protein staining

For imaging screen samples, after BARC-FISH imaging was completed, the nuclei were stained by applying 3 mL of 1:1000 diluted DAPI (Thermo Scientific, 62248) in 2×SSC in 4 minutes, and incubated for 3.5 minutes. The sample was then washed by applying 2 mL of wash buffer. Total protein stain was then conducted by diluting CF568-labeled succinimidyl ester (Biotium, 92131) at 1:100,000 in water containing 0.1 M NaHCO_3_ and 25 mM Na_2_CO_3_, flowing 3 mL of the buffer through the sample chamber over 4 minutes, and incubating for 3.5 minutes. After further application of 2 mL wash buffer and 2 mL imaging buffer, three z image stacks were taken at each FOV sequentially with 546-nm, 477-nm and 405-nm lasers. Specifically, the z-stack of 546-nm and 477-nm had the same step size, exposure time and total range as that used in BARC-FISH imaging, and the z-stack of 405-nm lasers had the same parameters as those used in chromatin tracing imaging. For imaging siRNA validation samples, DAPI staining was applied and imaged after chromatin tracing in a similar manner in the 477-nm and 405-nm channels, with parameters identical to those mentioned above.

#### siRNA knockdown

##### siRNA in A549 cells

The clonal A549-Cas9 cells used to construct the screen cell library were seeded at 6% density one day before siRNA transfection. The siCHD7 single knockdown experiments were conducted using Dharmacon ON-TARGETplus siRNA pools and DharmaFECT transfection reagent (Dharmacon, T-2001-03) following the manufacturer’s protocol. Briefly, for each 10-cm dish, 3 µL of 100 µM siCtrl (Dharmacon, D-001810-10) or siCHD7 (Dharmacon, L-025947-01) was diluted to 1.2 mL Opti-MEM reduced-serum medium (Thermo Fisher Scientific, 31985-070). In a separate tube, 24 µL of transfection reagent was diluted to 1.2 mL Opti-MEM reduced-serum medium. The diluted siRNA and the diluted transfection reagent were incubated for 5 minutes at room temperature. Then the diluted siRNA was added into the diluted transfection reagent and the mixture was incubated for 20 minutes at room temperature. Lastly, the antibiotics-free growth medium was added to a final volume of 12 mL. Cells were cultured with the 12 mL transfection medium at 37°C for 96 hours and used for follow-up analysis, including Western blot, chromatin tracing, RNA-seq, and CUT&RUN.

##### siRNA in hTERT-RPE1 cells

The hTERT-RPE1 cells were seeded at 6% density one day before siRNA transfection. The same siRNA pools for siCtrl and siCHD7 in A549 RNAi experiments and RNAiMAX transfection reagents (Invitrogen, 13778-075) were used. Transfection was done following RNAiMAX manufecturer’s protocol with 50nM final concentration of siCtrl or siCHD7 siRNA. After siRNA transfection, cells were cultured with the transfection mix for 4 days before downstream assays.

#### HDAC inhibition drug treatment

A549 cells were seeded at 40% density one day prior to the drug treatment. Trichostatin A (TSA) (Sigma-Aldrich, T8552) was dissolved in DMSO at 1mM to make drug stocks. To inhibit HDAC, cells were cultured in media supplemented with 150 nM TSA or equivalent volume of DMSO (as a solvent-only control) for 2 hours before PFA fixation.

#### Western blot

Cells were trypsinized by TrypLE Express (Thermo Fisher Scientific, 12605-010) and pelleted through centrifugation at 500g for 5 minutes at 4°C. To enrich the nuclear proteins, cell nuclei were isolated and lysed using NE-PER Nuclear and Cytoplasmic Extraction Kit (Thermo Fisher Scientific, 78833) following the manufacturer’s instructions. For hESCs and human neural crest progenitors, protein was extracted using RIPA buffer (Thermo Scientific 89900). The reagents were supplemented with 1× protease inhibitors (Thermo Fisher Scientific, 87786) to prevent the protein degradation during extraction. The protein concentrations were quantified using Pierce BCA protein assay kit (ThermoFisher, 23225) following the manufacturer’s instructions. The protein was denatured at 95-100°C for 5 minutes in 1× NuPAGE LDS sample buffer (ThermoFisher, NP0007) containing 10% v/v β-Mercaptoethanol (Millipore Sigma, M6250). For detection of CHD7 protein, the denatured nuclear extract was subjected to electrophoresis on a precast 3-8% Tris-Acetate polyacrylamide gel (Thermo Fisher Scientific, EA0378BOX) in 1× Tris-Acetate SDS running buffer (Thermo Fisher Scientific, LA0041) at 100V for 2.5 hours. For detection of CTCF and Actin, the protein electrophoresis was performed using 4-15% precast polyacrylamide gels (Biorad, 4568084) in 1× Tris/Glycine/SDS buffer (diluted from Biorad, 1610732) at 100V for 1.5 hours. For hESCs and human neural crest progenitors, the protein electrophoresis was performed using 4-20% precast polyacrylamide gels (Biorad, 4568095) in 1× Tris/Glycine/SDS buffer (diluted from Biorad, 1610732) at 100V for 2.5 hours for all proteins. The proteins were transferred from the gel onto a nitrocellulose membrane (ThermoFisher, IB301032) using either iBlot2 dry blotting system (Thermo Fisher Scientific, IB21001S) or wet transfer using the Biorad Mini Trans-Blot Cell following the manufacturer’s instructions. Wet transfer buffer was 20% methanol, 200 mM glycine, and 250 mM Tris. The membrane was blocked in 5% BSA in TBST for 1 hour at room temperature with gentle shaking. Primary antibodies against human CHD7 (Thermo Fisher Scientific, PA5-72964), CTCF (Millipore Sigma, 07-729), HSP90 (CST 4874S), Sox10 (CST 89356S) and Actin (Abcam, ab179467) were incubated with the membrane at recommended concentrations at 4℃ overnight with gentle shaking. The following day, the membrane was washed three times in TBST for 5 minutes each, followed by incubation with horseradish peroxidase-conjugated secondary antibodies (Abcam, ab6721) diluted at 1:3000 ratio in 5% non-fat milk in TBST for 1 hour at room temperature. The blots were then washed three times in TBST for 5 minutes each. CTCF and Actin blots were treated with SuperSignal West Pico PLUS Chemiluminescent Substrate (Thermo Fisher Scientific, 34580) and CHD7 blot was treated with SuperSignal West Femto Maximum Sensitivity Substrate (ThermoFisher, 34094) for 2-5 minutes at room temperature. The blots were imaged using a CCD camera-based imager, ProteinSimple FluorChem E system or Bio-Rad ChemiDoc MP Imaging System.

#### Immunofluorescence of Cas9 and H3K27ac

Cells were fixed in 4% PFA in DPBS for 10 minutes and washed twice with DPBS. Cells were then permeabilized in 0.5% v/v Triton X-100 in DPBS for 10 minutes and washed twice with DPBS. Cells were blocked in blocking buffer (1% w/v BSA and 0.1% v/v Tween-20 in DPBS) for 30 minutes at room temperature with gentle agitation. Then the cells were incubated for 1 hour at room temperature with 1:500 diluted anti-Cas9 primary antibody (Sigma, SAB4200701-25UL, for detection of Cas9 expression) or 1:100 diluted anti-H3K27ac primary antibody (Active Motif, 91193, for detection of H3K27ac level in TSA- and DMSO-treated samples) in blocking buffer, followed by three DPBS washes for 5 minutes each. The cells were incubated with 1:1000 diluted Alexa Fluor 647-labeled secondary antibody (Thermo Fisher Scientific, A21237, for anti-Cas9 primary antibody) or Alexa Fluor 488-labeled secondary antibody (Invitrogen, A21202, for anti-H3K27ac primary antibody) in blocking buffer for 1 hour at room temperature, followed by three DPBS washes for 5 minutes each. Starting from incubation with secondary antibody, the samples were covered from the light with aluminum foil.

#### Data analysis

##### Determination of chromatin traces in 3D

Image and phenotype analyses were performed using MATLAB version R2020a. We adopted a previously reported pipeline to determine the 3D chromatin traces^90^. Briefly, to correct the drift between sequential chromatin tracing secondary hybridization rounds, we determined the 3D positions of fiducial beads during each round of hybridization by fitting their z-stack images to Gaussian functions in 3D. We then fitted the center position of each DNA FISH foci in 3D using the same algorithm, and the sample drift (represented by bead movements) was subtracted from the 3D position of each DNA locus. The drift-corrected DNA loci positions were then linked into traces based on their expected spatial proximity in each chromosome territory. To remove false traces generated from non-specific target objects or genomic DNA released from dead cells, linked traces were excluded if 1) they located outside of the nucleus, represented by 2D-projected DAPI staining patterns, or 2) their radii of gyration was smaller than 0.5 μm. In a small number of datasets (see *Chromatin tracing sequential imaging* section for explanation), we noticed that due to extra-strong labeling of TAD4, signal from TAD4 (638-nm channel) could be excited with 546-nm laser and thus be misidentified as TAD18. To correct this in the affected datasets, after the traces were linked, we calculated the 3D distance between TAD4 and TAD18 on each trace in those datasets. If the distance was smaller than 100 nm, the 3D position recognized as TAD18 were regarded as bleedthrough from TAD4, and the 3D positions of TAD18 were assigned to NaN.

##### BARC-FISH pattern extraction

The decoding process consisted of the following steps: 1) drift correction, 2) identifying all possible BARC-FISH patterns for each value of each barcode digit, 3) filtering mis-identified BARC-FISH patterns based on intensity thresholding, and 4) assigning the digit identity to each cell. To correct the drift between sequential BARC-FISH imaging rounds, the center positions of the fiducial beads were determined by fitting to Gaussian functions in 2D, and the movement was subtracted from BARC-FISH images. To extract the BARC-FISH patterns for each value in each digit, BARC-FISH images were first projected along z dimension with average intensity projection, and filtered by a 2D median filter and a Gaussian low-pass filter. The image background was derived by image opening with a disk-shaped structural element with a 20-pixel radius, and the background was subtracted. Potential BARC-FISH areas were identified by binarizing the background-subtracted image and selecting objects that were larger than 50 pixels. Due to the strong labeling intensity of BARC-FISH foci, bleedthrough was often seen in channels excited with lasers of shorter wavelength. To correct this, the bleedthrough of Alexa Fluor 647 into 546-nm channel was identified by the shared areas between 546-nm and 638-nm channel in the same BARC-FISH round, and the areas were subtracted from the 546-nm BARC-FISH pattern. Similarly, the bleedthrough of Alexa Fluor 750 into 638-nm channel was subtracted from the 638-nm BARC-FISH pattern. Regions that were positive for all three channels (546-nm, 638-nm and 749-nm) during one BARC-FISH round were subtracted from all three channels because such regions could be due to broad-spectrum autofluorescent objects or non-specific binding of all probes to the same sticky objects. These steps generated high-fidelity binary masks for BARC-FISH signals. To further increase the reliability of our identified BARC-FISH foci masks, we applied the binary masks to the median- and Gaussian-filtered BARC-FISH images collected in 15-20% of the FOVs collected in each screen dataset. For each value of each digit, we calculated the normalized intensity ([total intensity] / [pattern size]) of each BARC-FISH focus identified through the binary mask, and sorted the normalized intensity. A normalized intensity cutoff was then derived for each value of each digit, and applied to all the FOVs in the dataset to only include the BARC-FISH patterns of which the normalized intensity was above this threshold.

##### Cell segmentation and decoding

Drift between the DAPI/total protein stain imaging round and previous rounds were corrected using the same algorithm as in BARC-FISH pattern extraction. Z-stacks of DAPI and total protein stain was projected along the z dimension with max and average intensity projections, respectively. A median filter was applied to the z-projected DAPI images. To remove the bleedthrough of the fiducial beads pattern into total protein stain, total protein images was eroded by a disk-shaped structural element with a radius of 5 pixels. We then performed watershed-based cell segmentation of the total protein stain profile using DAPI patterns as foreground markers. To determine the barcode identity of each cell, for each digit, we calculated the total pixel counts from BARC-FISH pattern of each of the three values, value 0 (546-nm channel), 1 (638-nm channel) or 2 (749-nm channel), and assigned the value with the largest pixel count to the digit of the cell. Cells with BARC-FISH signal in none of the 3 values were assigned a fourth value of 10000. We then calculated the Hamming distance between the current decoded barcode to those in the codebook. A barcode was decoded/corrected if: 1) the current decoded barcode has a unique closest match in the codebook, 2) this unique closest match is a good code (a code that uniquely associates with only one sgRNA), and 3) the Hamming distance between the decoded barcode and the closest match was not larger than 1 (maximum 1 mismatching digit allowed). The cell carrying the decoded/corrected barcode was assigned the corresponding sgRNA identity.

##### Cell cycle stage classification

We only included G1-phase cells in phenotypic analyses of both the screen and the validation. G1 cells were indicated by an absence or low intensity of Geminin stain in the nucleus. The foreground DAPI marker generated for the watershed-based cell segmentation was used as a binary mask of nuclear regions. Two methods were used to determine the cell cycle status: 1) We calculated the normalized Geminin intensity for each cell by dividing the total nuclear Geminin fluorescence intensity by the area of the nucleus, and empirically determined the threshold for each imaging experiment between Geminin-positive and Geminin-negative cells based on the distribution of the normalized intensity. Cells of which the normalized nuclear Geminin intensity was below the threshold were included in phenotypic analyses. This method was used for 13 out of 17 screen experiments and all validation experiments. For 4 early replicates, due to area-to-area variation of Geminin staining, a uniform normalized nuclear Geminin intensity could not be applied to all FOVs for G1 cell identification. For these 4 replicates, we determined the cutoff for G1 versus non-G1 cells by calculating the ratio between the normalized Geminin intensity in the nucleus and the whole cell area (determined by watershed boundaries) for each cell, and empirically determined the threshold for each dataset based on the distribution of this ratio.

##### Analysis of adjacent TAD distance and contact

Supplementary Table 13 includes the exact sample size (*n*), the p-value, false discovery rate (FDR) and log2 fold change (log2fc) value for each analyzed sgRNA and for each phenotype analyzed in the screen. We grouped traces from the same sgRNA (determined by the cell barcode detected via BARC-FISH) across different datasets. To increase the statistical power in identifying novel hits, for phenotype analyses regarding chromatin folding and nuclear lamina association, traces from all 412 observed sgRNAs in the screen were pooled as control (termed as “whole population”), and 341 sgRNAs with at least 40 chromatin traces were compared to the whole population, which were used to generate volcano plots (Fig. 2a,d,f,h,j, and Fig. 5a) and Fig. 2b. In other panels, only non-targeting control sgRNAs with no fewer than 40 traces were pooled and plotted as sgCtrl, and statistics were performed between selected hits and the included non-targeting control sgRNAs (Fig. 2c,e,g,i,k,l, Fig. 3 and Fig. 5b,c). Adjacent TADs were defined as TADs that were next to each other along the genomic map. To compare adjacent TAD distance, we calculated the distance between each pair of adjacent TADs on each trace, and generated a mean adjacent TAD distance profile for each sgRNA by calculating the mean distance for each pair of adjacent TADs. We then conducted two-sided Wilcoxon signed rank test in MATLAB to compare the mean adjacent TAD distance between each sgRNA and the control group, calculated FDR from the resulting p values, and selected top hits that 1) have FDRs<0.1, 2) have the largest absolute values of log2fc, and 3) have protein localization in the cell nucleus based on The Human Protein Atlas (proteinatlas.org) to prioritize potential direct regulators of 3D genome. To calculate the log2 fold change in Fig. 2a, we divided the mean adjacent TAD distance between the same pair of TADs from the sgRNA by that from the control group, calculated the log2 fold change value for each TAD pair, and averaged the log2 fold change values to generate the log2 fold change for each sgRNA. Log2 fold change was calculated in a similar manner in Fig. 2j by averaging the log2 fold change value of all pairwise comparisons. To analyze adjacent TAD contact frequency (Extended Data Fig. 8b,c), we defined a contact event as when the spatial distance between two TADs is less than 500 nm, and counted the frequency of contacts between each pair of adjacent TADs.

##### Determination of compartment identity of TADs

We assigned A/B compartment identity to the TADs using a previously introduced algorithm^90^. Briefly, we fitted a power-law function to the data points of the mean inter-TAD spatial distances versus their genomic distances, which yielded the expected inter-TAD spatial distance for each pair of TADs according to their genomic distance. We then normalized the observed mean inter-TAD spatial distance by the expected spatial distance, yielding a normalized inter-TAD distance matrix. We then calculated the Pearson correlation coefficient between each pair of rows/columns in this matrix, generating a Pearson correlation matrix. We next applied principal component analysis to the Pearson correlation matrix, and took the coefficients of the first principal component as compartment scores. The compartment score profiles were cross-validated with published ChIP-seq profiles^96, 97^, so that the compartment A regions are enriched in active histone modifications, while compartment B regions are enriched in repressive histone modifications. If the trend was opposite, the signs of the compartment scores were flipped. In the CRISPR screen, traces from the whole population were pooled to generate the compartment identity, which was applied to different perturbations when evaluating long-range AA, AB and BB compartment contacts (Fig. 2d-i).

##### Analysis of long-range contact in different compartments

Long-range contacts were defined as the spatial contact events between TAD pairs that are not adjacent TADs on the genomic map. We used the same distance threshold for calling contacts as that used in the adjacent TAD contact, and calculated the numbers of AA, AB and BB long-range TAD contact events per trace. In Fig. 2d-i and Fig. 3, AB compartment profile generated from all traces were used to define A and B compartments. In Extended Data Fig. 3e, 4d, and 5e, AB compartment profile of the according control group (siCtrl of A549-Cas9 cells, GFP-overexpressed A549-Cas9 cells and siCtrl of hTERT-RPE1, respectively) were used to define A and B compartments.

##### Analysis of overall chromosome compaction

We calculated the inter-TAD distances between all pairs of TADs on chr22 for each sgRNA, and generated mean inter-TAD distance matrix for each sgRNA by calculating the mean distance between each pair of TADs. We then performed two-sided Wilcoxon signed rank test between the mean inter-TAD distances of each sgRNA and control, and calculated FDR and log2 fold change values.

##### Determination of chromatin association with nuclear lamina

The 2D-projected nuclear lamina profile was derived from 2D-projected DAPI images by adopting a previously reported algorithm^90^. Briefly, we extracted the pixels on the edges of average z-projected nuclei and measured the x-y 2D distance between the positions on each trace (trace centroid or each TAD position) and the nearest nuclear edge pixels. The trace centroid distances to nuclear lamina of each sgRNA and control were compared.

##### Analysis of nuclear morphological properties

To increase the statistical power of the screen, similar to analyzing chromatin phenotypes, 3D nuclear DAPI patterns from all 412 sgRNAs were pooled as controls in the volcano plots (Fig. 5d,h), and only sgRNAs with more than 10 nuclei were included in the screen analyses; 3D nuclear DAPI patterns from non-targeting control sgRNAs with more than 10 nuclei were pooled as controls in the box plots (Fig. 5e-g,i,j). We compared the 3D coefficient of variation of nuclear stain intensity (nuclear intensity unevenness) and nuclear sphericity between each sgRNA and control. To abstract the nucleus shape in 3D, we first calculated a normalization factor of each FOV by selecting the max intensity of the median-filtered z-stack images of the current FOV. We normalized the max z-projected DAPI image by this normalization factor, and calculated a background by applying adaptive thresholding on the normalized image using a sensitivity of 0.4. We then processed each layer of the z-stack, normalized the median-filtered current slice of z-stack image by the normalization factor, and binarized it using the threshold generated from the previous adaptive thresholding step. Only binary objects at each slice that were larger than 5,000 pixels were retained. Partial nuclei at the edges of each FOV were excluded. We then matched the cell identity to each segmented 3D nucleus, and analyzed the coefficient of variation of voxel intensity within each nucleus of each sgRNA. The nuclear volume and surface area were calculated by the *regionprops3* function in MATLAB. The nuclear sphericity was determined by the following equation:

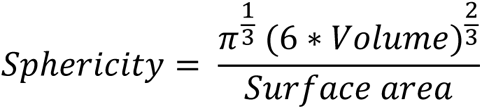

A sphericity of 1 indicates the shape is a perfect sphere. A sphericity smaller than 1 indicates reduced roundness. Sphericity values exceeding 1 due to inaccuracy of surface area calculation by the *regionprops3* function were excluded from the analysis. For volcano plots, p values were calculated by unpaired t test followed by FDR calculation (Fig. 5d,h). For box plots, p values were calculated by two-sided Wilcoxon rank sum test (Fig. 5e,i).

##### Calculation of radius of gyration

We only retained traces with more than 80% of TADs detected for analyzing this phenotype. The radius of gyration was defined as the root mean square of the distance between each TAD to the centroid of the trace, as shown in the following equation:

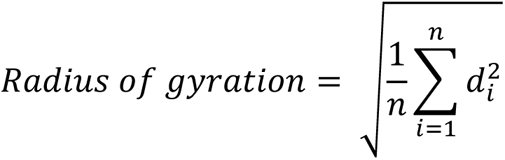

where *n* is the number of detected TADs in the given trace, and *d_i_* represents the spatial distance between TAD *i* and the centroid of the trace.

##### Quantification of polarized organization of A-B compartments

Only traces with more than 80% of TADs detected were included in this analysis. We used a previously developed algorithm^90^ to quantify the polarized organization of A-B compartments, yielding a polarization index. Briefly, we constructed 3D convex hulls for all A compartment TADs and B compartment TADs on each individual trace, with the *convhull* MATLAB function. We then calculated the volume of these two hulls (referred to as V_A_ and V_B_, respectively) and the volume of the overlapped spaces between A and B hulls (referred to as V_S_). Polarization index was defined as:

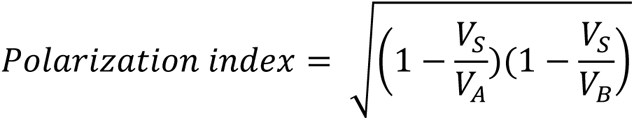

A polarization index of 1 indicates that the A and B compartments are completely spatially separated from each other in a side-by-side manner. A polarization index of 0 indicates that A and B compartments completely overlap, or one is surrounded by the other. The A-B compartment profile of the according treatment was used to calculate polarization indices for each condition in Extended Data Fig. 3d, 4c.

#### RNA-seq

##### Library preparation

Total RNA from siCtrl and siCHD7 A549-Cas9 cells was extracted using Qiagen RNeasy Mini Kit (Qiagen, 74104) with on-column DNase treatment (Qiagen, 79254) according to the manufacturer’s specifications. RNA integrity was determined using Agilent Fragment Analyzer RNA Kit (DNF-471) according to the manufacturer’s specifications. To prepare the sequencing library, for each sample, 1 μg of RNA was prepared using Kapa Biosystems KAPA mRNA HyperPrep kit with 10 PCR cycles according to the manufacturer’s specifications. The final library quality control was run on the Agilent TapeStation D1000 assay, and the concentration was determined with the KAPA Library Quantification Kit. Sequencing was performed with 25 million reads per sample, 100 bp paired-end reads, on an Illumina NovaSeq 6000. 4 replicates of siCtrl and 4 replicates of siCHD7 cells were processed in this study.

##### RNA-seq data analysis

RNA-seq reads from siCHD7 and siCtrl-treated A549-Cas9 cells were aligned to Ensembl genome GRCh38 by STAR version 2.7.9^98^. Counts of each unique gene were performed by FeatureCounts contained in the Subread package (2.0.0)^99^. Differentially-expressed genes (DEGs) were detected using the DESeq2 R package from Bioconductor^100^. Cutoffs were placed at > 20 counts per gene, Log2FC > 0.5, and P Value > 10e-10. The expression levels were normalized and adjusted using Apeglm^101^. Bioconductor EnhancedVolcano (http://bioconductor.org/packages/release/bioc/html/EnhancedVolcano.html) was used to generate the volcano plot and previously listed cutoffs were added to the plot. Ensembl IDs were converted to gene names using Bioconductor package biomaRt (version 2.28.0)^102^. Gene ontology (GO) enrichment analysis was performed using Enrichr GO:BP 2021^103^. Log10 P values were generated and listed on the graph with the top 10 up and down GO terms after analysis of all significantly upregulated genes and downregulated genes according to cutoffs.

#### Condensation assays

##### CHD7 purification protocol

Human CHD7 with N-terminal 6xHis tag and C-terminal Flag tag was purified as previously described with minor modifications^26^. Briefly, the pFastBac-CHD7 vector (Addgene, 170141) was transfected into DH10Bac E. coli to produce bacmids following Bac-to-Bac (Thermo Fischer Scientific, 10359016) instructions. The proteins were purified either from monolayer Sf9 cell cultures transfected with bacmids using ExpiFectamine (Thermo Fischer Scientific, A38915) or from Sf9 suspension cultures infected by P0 baculovirus. The infected cells were washed with PBS and resuspended in BC buffer (10% Glycerol, 20 mM HEPES, pH 7.9, 0.4 mM EDTA, supplemented with protease inhibitors (complete tablets, Roche)) containing 250 mM NaCl (BC250) as well as 5 mM imidazole. Cells were lysed by three freeze-thaw cycles in liquid nitrogen and 42°C water bath, followed by passing through an 18-gauge needle for three times to shear DNA. The lysates were then cleared by centrifugation before adding Ni-NTA resin (Invitrogen, R901) that has been equilibrized with BC250 buffer. The Ni-NTA resin was incubated with the lysate for 1 hour at 4°C, washed with 5x resin volume of BC250 buffer containing 10 mM imidazole for three times, and eluted with 5x resin volume of BC250 buffer containing 200mM imidazole. The elute was then concentrated using Amicon Ulrea-4 Ultracell-100K centrifugal filters (Millipore, 910024) and imidazole concentration was diluted down to 50 mM before adding Pierce™ Anti-DYKDDDDK Affinity Resin (Thermo Fischer Scientific, A36801) and incubating for 4 hours at 4°C. The resin was then washed sequentially with 10x resin volume of BC500 buffer twice, 10x resin volume of BC1000 buffer twice, 10x resin volume of BC500 buffer once, 10x resin volume of BC200 buffer once and 10x resin volume of BC100 buffer once. Double-tagged full-length CHD7 proteins were finally eluted using 10x resin volume of BC100 buffer containing 0.25 mg/mL FLAG-peptide (Sigma, F3290) after 1-hour incubation at 4°C. Purified CHD7 proteins were further dialyzed into 1x phasing buffer (20 mM HEPES pH 7.4, 70 mM KCl, and 1 mM DTT) at 4°C overnight and concentrated to around 1mg/mL using Amicon Ulrea-4 Ultracell-10K centrifugal filters (Millipore, 901024) before usage.

##### PEG Silane coating of 384-well plates

*In vitro* imaging was performed in 384-well plates (Sigma-Aldrich, M4437). The plates were coated with PEG Silane as previously described^104^. Briefly, individual wells were first washed with 100 µL of 2% Hellmanex (Sigma-Aldrich, Z805939) for 1 hour and rinsed three times with water. Next, each well was incubated with 0.5 M NaOH for 30 minutes and rinsed three times with water. Then, each well was incubated 100 µL of 20 mg/mL PEG-silane MW-5000 (Laysan Bio, MPEG-SIL-5000) dissolved in 95% EtOH at 4°C overnight. On the next day, the plate was first rinsed three times with water and incubated with 100 mg/mL BSA (Sigma-Aldrich, A9647) for 30min. Finally, each well was rinsed three times with water and 1x phasing buffer sequentially. 10 µL of 1x phasing buffer was maintained at the bottom of each well to prevent drying before usage.

##### Condensation assays

First, 150 ng/µL bacteriophage lambda-DNA (Thermo Fisher Scientific, SD0011) was pre-incubated with 5x GelGreen (Biotium, 41004) in 0.5x phasing buffer for dye labeling. Next, 10 µL of 3 µM CHD7 solution, 10 µL of 1x phasing buffer, and 1.5 µL of the dye-labeled lambda DNA solution were mixed in a 1.5 mL tube. 20 µL of 1x phasing buffer and 1.5 µL of the dye-labeled lambda DNA solution were mixed as negative control. For each condition, 20 µL of mixture solution were pipetted into a well of the PEG Silane coated 384-well plate, and condensates were imaged using a Nikon Eclipse TS100 Inverted Routine Microscope (NI-TS100) with Nikon CFI Plan Fluor 4X Objective and Nikon C-HGFI Intensilight Epi-fluorescence Illuminator.

#### CUT&RUN

##### CUT&RUN protocol

CUT&RUN was performed using the Epicypher CUT&RUN Kit (14-1048) according to the manufacturer’s specifications with the following conditions and modifications: For binding to the Concanavalin beads, 500,000 cells per sample were prepared, counted twice by hemacytometer and averaged, and incubated with activated beads. Antibodies were used as listed below, with 0.5 ug of antibody per reaction.

H2A antibody – CST Histone H2A (D6O3A) Rabbit mAb #12349 IgG Control antibody – CUTANA Kit Rabbit IgG CUT&RUN Negative Control Antibody H3K4me3 antibody – CST Tri-Methyl-Histone H3 (Lys4) (C42D8) Rabbit mAb #9751 CTCF antibody – CTCF CUTANA™ CUT&RUN Antibody 13-2014 CHD7 antibody – Invitrogen CHD7 Polyclonal Antibody PA5-72964 RAD21 antibody – Active Motif Rad21 #91245 *Library preparation.* Libraries were prepared using the Epicypher CUT&RUN Library Prep Kit (14-1001) according to the manufacturer’s specification with the following modification: SPRI-select beads were used after library preparation to perform right and left-handed selections at size 200 bp – 700 bp to enrich for DNA fragments from CUT&RUN and remove any adapter dimers or high molecular weight DNA. Library quality was analyzed using Agilent Tapestation D1000 High Sensitivity Tapes #5067-5584. Sequencing was performed with 150 bp paired-end reads, on an Illumina NovaSeq 6000.

##### CUT&RUN data analysis

CUT&RUN reads were analyzed for quality using FastQC (https://www.bioinformatics.babraham.ac.uk/projects/fastqc/) and trimmed using Trimmomatic^105^. Reads were then aligned to hg18 by Bowtie2 version 2.3.4^106^. SAMTools version 1.11 was then used to convert to BAM format, index, isolate uniquely mapped paired reads, and remove duplicates^107^. Picard version 2.27.4 was used to downsample all samples to the same read depth (5 million reads, with CHD7 at 12 million reads) for direct comparison. MACS2 version 2.2.7.1 was used to call sample narrow peaks using IgG as an input. IgG was always downsampled to the exact depth of the sample it was serving as input for. Read counts across genomic intervals and peak visualization were performed using deepTools version 3.3^108^, and .bw files were visualized using Integrative Genomics Viewer.

#### RNA MERFISH

##### MERFISH probe design

Sequences in the MERFISH template oligo library were derived from those reported in a previous report^66^ with the following modifications: 1) the priming regions for primary probe amplification were replaced; 2) the rest of the sequences were reverse-complemented, since our *in vitro* transcription and reverse-transcription ran in the opposite directions to those in the previous report (Supplementary Table 14). The template library was ordered from GenScript.

##### MERFISH primary probe hybridization

A549-Cas9 cells treated with siCtrl and siCHD7 were labeled with geminin following the previously described procedure, *Geminin antibody staining*. After post-fixation, cells were incubated in pre-hybridization buffer (50% formamide in 2×SSC) for 5 minutes at room temperature. Then the cells were incubated with hybridization buffer (50% formamide, 0.1% w/v yeast tRNA, 10% v/v dextran sulfate, 1% v/v murine RNase inhibitor and 4 μM primary probes in 2×SSC) at 37℃ for 16-20 hours. Cells were washed with 2×SSCT (0.1% v/v Tween-20 in 2×SSC) twice in a 60°C water bath and once at room temperature, for 15 minutes for each wash. The yellow-green fiducial beads were applied to the cells at 1:250,000 dilution in 2×SSC and incubated for 10 minutes at room temperature. The unattached beads were removed by two brief washes with 2×SSC. The coverslips were assembled in the FCS2 flow chamber and mounted onto the microscope for sequential hybridization and imaging.

##### MERFISH sequential hybridization and imaging

We adapted the MERFISH sequential imaging protocol in a previous report^90^ using the same imaging system described previously in *Imaging system setup*. We conducted 8 rounds of three-color imaging in a similar manner to *Chromatin tracing sequential imaging* to detect MERFISH signals and fiducial beads. During each round of hybridization, the sample was incubated with two secondary probes (one labeled by Alexa Fluor 647 and one labeled by Alexa Fluor 750) diluted in 20% v/v EC, 2×SSC containing 1:1000 diluted MRI for 25 minutes. The final concentration of each secondary probe was 3 nM. At each FOV, three z image stacks were taken sequentially with 750-nm, 647-nm and 477-nm laser illumination, with a step size of 500 nm, 0.4 s exposure time at each step, and a total z range of 7.5 μm. After all the FOVs were imaged for the current imaging round, the secondary probes were stripped off by slowly flowing 4 mL of 65% v/v formamide in 2×SSC for 10 minutes and incubating for additional 5 minutes. After all 8 rounds of MERFISH imaging was finished, 1:1000 diluted DAPI in 2×SSC was applied to the sample, let stand for 2 minutes, and excessive DAPI was washed off by applying 2×SSC. Two z image stacks were taken sequentially with 477-nm and 405-nm laser illumination, with a step size of 500 nm, 0.4 s exposure time at each step, and a total z range of 7.5 μm to image DAPI stain. The secondary probes were ordered from IDT and their sequences are listed in Supplementary Table 15.

##### MERFISH analysis

We adopted a previously reported pixel based MERFISH decoding^66^. First, to correct the drift between MERFISH hybridization rounds, we determined the 2D positions of fiducial beads by 2D Gaussian fitting for each round of hybridization. A transformation matrix file for each field of view in each round was generated and used for image registration and correction of x-y drift. Next, for each z-stack image, the background was removed by subtracting an image generated by image opening with a disk-shaped morphological structuring element with a radius of five pixels. To decode the MERFISH foci, the values of each pixel in 16 rounds were concatenated into a 16-dimensional “pixel vector”. The pixel vectors were normalized to generate unit vectors and compared with unit vectors of all 140 correct codes in the codebook (Supplementary Table 16). The closest correct code for each pixel was identified if it is within a code distance (in the 16-dimensional space) error threshold of 0.6058. The adjacent pixels matched to the same code were combined as one molecule. To further select real RNA molecules, we created a parameter space including three properties: the foci area, the distance to the matched code, the magnitude of the pixel vector. The parameter space was segmented into small bins and the error rate of each bin was calculated by the ratio of foci number that matched to empty control codes over foci number that matched to all 140 codes. The foci with parameters in bins with an error rate smaller than a cutoff threshold will be finally selected. For each dataset, 20 random field of views were used to define the parameter space and an adaptive procedure was performed to determine the cutoff threshold so that the overall error rate is below 5%. Finally, the first round of MERFISH signal were used for watershed-based cell segmentation and decoded RNA foci were assigned to each cell. Only G1 cells were selected for downstream analysis by geminin stain signals.

#### Poly-A stain

The siCtrl- and siCHD7-treated A549-Cas9 cells were stained with geminin following the procedure of *Geminin antibody staining*. Then, cells were incubated with pre-hybridization buffer containing 50% formamide in 2×SSC for 5 minutes at room temperature. The hybridization buffer containing 50% formamide, 0.1% w/v yeast tRNA, 10% v/v dextran sulfate, 1% v/v murine RNase inhibitor and 2 μM poly-dT adapter probe (/5Acryd/TTGAGTGGATGGAGTGTAATT+TT+TT+TT+TT+TT+TT+TT+TT+TT+T) in 2×SSC was applied to the cells and incubated at 37℃ for 16-20 hours. The excessive probes were removed by two washes with 2×SSCT in a 60°C water bath, and once with 2×SSCT at room temperature for 15 minutes each. The Alexa Fluor 750 dye-labeled readout probe (/5Alex750N/ATTACACTCCATCCACTCAA) was incubated with the cells at 10 nM in 20% v/v EC, 0.1% v/v murine RNase inhibitor in 2×SSC for 30 minutes at room temperature, followed by two washes with 20% v/v EC in 2×SSC for 5 minutes each at room temperature to remove the unbound readout probes. The cells were stained with DAPI at 1:1,000 in 2×SSC for 5 minutes at room temperature, and rinsed twice with 2×SSC. The cells were submerged in oxygen-scavenging imaging buffer that contained 50 mM Tris-HCl pH 8.0, 5% w/v glucose, 2 mM Trolox, 0.5 mg/mL glucose oxidase, 40 μg/mL catalase and 0.1% v/v murine RNase inhibitor in 2×SSC for imaging. Cells were imaged in 749-nm channel for poly-A stain, 477-nm channel for geminin stain, and 405-nm channel for DAPI stain using the same microscope setup described previously. The z-stack images were taken with a step size of 200 nm, 0.4 s exposure time at each step, and a total z range of 9 μm for poly-A and DAPI stain, and 7 μm for geminin stain. To analyze the images, we first extracted the nuclear area by taking the mean projection of the z-stack images for DAPI. We then applied the DAPI mask to the mean projection of z-stack images for poly-A, summed the intensity values within each nucleus, and divided the sum by the area of each nucleus. Only G1 cells were included for this analysis by geminin stain signals.

#### Chromatin polymer simulation

A Monte Carlo simulation of chromatin conformation with a lattice polymer model was adapted from a previous report^90^ with the following modifications: 1) The polymer contained 30 monomers; 2) the maximum allowed distance between adjacent monomers along the polymer was 3; 3) to calculate the energy of a polymer conformation, we included only one type of interaction energy – any pair of monomers with a distance closer than 2 would incur an energy loss of K. To build a bounding envelope of the simulated chromatin conformation, at each monomer, we created a sphere with a radius of 15 pixels, the centroid of which is the current monomer. We then filled and smoothened the “dilated” 3D shape by function *imfill* and *imclose* in MATLAB. We generated the isosurface of the dilated shape by MATLAB function *isosurface* to define the bounding envelope of the chromatin conformation for sphericity calculation and visualization in Fig 6d.

**Supplementary Table 1 (Separate file).**

Target gene summary. Three categories of target genes are included in the screen, as described in the table.

**Supplementary Table 2 (Separate file).**

Disease relevance of top hits identified in the screen.

**Supplementary Table 3 (Separate file).**

Digits and values for DNA barcode sequences in BARC-FISH screen, the corresponding linear and padlock probe sequences, and dye-labeled secondary probes.

**Supplementary Table 4 (Separate file).**

Oligonucleotide sequences of helper probes.

**Supplementary Table 5 (Separate file).**

Oligonucleotide sequences and PCR primers for barcode library assembly.

**Supplementary Table 6 (Separate file).**

Protospacer sequences for gene targets in the screen, and the oligonucleotide sequences and PCR primers for sgRNA library assembly.

**Supplementary Table 7 (Separate file).**

PCR primers for sgRNA-barcode CRISPR screen library cloning.

**Supplementary Table 8 (Separate file).**

Primers for CHD7 overexpression plasmid cloning.

**Supplementary Table 9 (Separate file).**

Oligonucleotide sequences for sgRNA-barcode sequencing library preparation.

**Supplementary Table 10 (Separate file).**

BARC-FISH codebook determined by sequencing. The “Goodcodes” worksheet contains good barcodes that each uniquely associate with an sgRNA, in which the first column represents the 10-digit barcode, and the second column represents the name of the sgRNAs listed in Supplementary Table 6. The “Badcodes” worksheet contains barcodes that associate with multiple sgRNAs.

**Supplementary Table 11 (Separate file).**

Template oligonucleotide sequences, PCR primers and reverse transcription primer used for chromatin tracing primary probe library synthesis.

**Supplementary Table 12 (Separate file).**

Secondary probe sequences for chromatin tracing.

**Supplementary Table 13 (Separate file).**

BARC-FISH screen results summary. 8 worksheets regarding 8 phenotype analyses were included, and each worksheet consists of the following columns:

sgRNA id: sgRNA ranking alphabetically, representing their orders in Supplementary Table 6;
sgRNA name: name of the sgRNAs as represented in Supplementary Table 6;
copy number/cell number: number of traces (for chromatin-related phenotypes) or cells (for nuclear morphological properties) analyzed;
p, fdr, log2fc: p value, false discovery rate, and log2 fold change of the analyzed phenotypes.

**Supplementary Table 14 (Separate file).**

Template oligonucleotide sequences, PCR primers and reverse transcription primer used for MERFISH primary probe library synthesis.

**Supplementary Table 15 (Separate file).**

Secondary probe sequences for MERFISH.

**Supplementary Table 16 (Separate file)**

Codebook for MERFISH.

**Supplementary File S1 (Separate file).**

Backbone sequence used for barcode plasmid library construction.

**Supplementary File S2 (Separate file).**

Example barcode plasmid sequence.

**Supplementary File S3 (Separate file).**

Backbone sequence used for sgRNA plasmid library construction.

**Supplementary File S4 (Separate file).**

Example sgRNA plasmid sequence.

**Supplementary File S5 (Separate file).**

Backbone sequence used for CRISPR screen plasmid library construction.

**Supplementary File S6 (Separate file).**

Example CRISPR screen plasmid sequence.

**Supplementary File S7 (Separate file).**

Sequence of plasmid for CHD7 overexpression.

**Supplementary File S8 (Separate file).**

Sequence of plasmid for CHD7-ΔBRK overexpression.

**Supplementary File S9 (Separate file).**

Sequence of plasmid for CHD7-K999R overexpression.

